# Neuronal lipolysis participates in PUFA-mediated neural function and neurodegeneration

**DOI:** 10.1101/2020.02.13.948430

**Authors:** Leilei Yang, Jingjing Liang, Sin Man Lam, Ahmet Yavuz, Meng C. Wang, Guanghou Shui, Mei Ding, Xun Huang

## Abstract

Lipid droplets (LDs) are dynamic cytoplasmic organelles present in most eukaryotic cells. The appearance of LDs in neurons is not usually observed under physiological conditions, but is associated with neural diseases. It remains unclear how LD dynamics is regulated in neurons and how the appearance of LDs affects neuronal functions. We discovered that mutations of two key lipolysis genes *atgl-1* and *lid-1* lead to LD appearance in neurons of *Caenorhabditis elegans*. This neuronal lipid accumulation protects neurons from hyperactivation-triggered neurodegeneration, with a mild decrease in touch sensation. We also discovered that reduced biosynthesis of polyunsaturated fatty acids (PUFAs) causes similar effects, synergistically with decreased lipolysis. Furthermore, we demonstrated that these changes in lipolysis and PUFA biosynthesis increase PUFA partitioning toward triacylglycerol, and reduced incorporation of PUFAs into phospholipids increases neuronal protection. Together, these results suggest the crucial role of neuronal lipolysis in regulating neural functions and neurodegeneration cell-autonomously.

**Highlights:**

1. Neuronal lipolysis prevents LD accumulation in neurons.
2. Defective neuronal lipolysis leads to touch sensation defect.
3. Blocking neuronal lipolysis alleviates neurodegeneration.
4. Neuronal lipolysis and *de novo* PUFA biosynthesis have a synergistic effect in neurodegeneration.
5. The incorporation of PUFAs into phospholipids promotes neurodegeneration.

## Introduction

Lipid droplets (LDs) are dynamic cytoplasmic organelles which are present in most, if not all, eukaryotic cells and many prokaryotic cells. By storing excess lipids in the form of neutral lipids including triacylglycerol (TAG) and sterol ester (SE), LDs maintain cellular lipid homeostasis, along with the coordinated actions of lipogenesis and lipolysis (Chen, Chen et al., 2019, Olzmann & Carvalho, 2019). The nervous system, including neurons and glia, has a high concentration of lipids. However, LDs in the nervous system are generally found in glial cells but not in neurons under normal conditions *in vivo* (Kis, Barti et al., 2015).

The origin and the role of glial LDs have been investigated only in recent years. The formation of LDs in glia, which act as a niche for neuroblasts, preserves *Drosophila* larval neuroblast proliferation under ROS-inducing stress conditions such as hypoxia. It is postulated that the incorporation of polyunsaturated fatty acids (PUFAs) into neutral lipids, and their storage in LDs, reduces the ROS insult and the toxic peroxidation of PUFAs in neuroblasts (Bailey, Koster et al., 2015). Several other studies reported the neuronal origin of glial LDs. When *Drosophila* neurons are under ROS insult or have mitochondrial dysfunction, neuronal lipid production is increased through SREBP-mediated lipogenesis. Interestingly, instead of forming LDs in neurons, lipids are transferred to neighboring glia through fatty acid transfer protein (FATP) or apolipoprotein to form LDs (Liu, MacKenzie et al., 2017, Liu, Zhang et al., 2015). In cultured hippocampal neurons, hyperactivated neurons also produce excess fatty acids, which are transferred, via lipid particles associated with ApoE, to astrocytes and are incorporated into LDs. The storage of fatty acids in astrocyte LDs and their subsequent β-oxidation in mitochondria protects neurons during periods of enhanced activity (Ioannou, Jackson et al., 2019). These findings suggest that LD formation in glia plays a role in protecting neurons from stress insults. However, it is unknown why neurons do not form LDs in an autonomous fashion to protect themselves under stress conditions.

Although neurons do not normally have LDs, some neuronal diseases are associated with LD biology and neuronal LDs have been reported in some disease models. The Parkinson’s disease protein α-Synuclein is located on the surface of LDs and α-Synuclein expression is correlated with LD accumulation (Cole, Murphy et al., 2002, Outeiro & Lindquist, 2003). Huntington’s disease cells, including primary striatal neurons and glia in Huntington’s disease mice, have dramatically increased LDs (Martinez-Vicente, Talloczy et al., 2010). Several hereditary spastic paraplegia (HSP) proteins affect LD dynamics, such as Spartin, spastin, atlastin-1, seipin and REEP1 (Ding, Yang et al., 2018, Eastman, Yassaee et al., 2009, Ebihara, Ebihara et al., 2015, Klemm, Norton et al., 2013, Papadopoulos, Orso et al., 2015, Renvoise, Malone et al., 2016). Despite the apparent association between LD and neuronal diseases, the causal link between neuronal LD dynamics and neuronal disorders remains largely elusive.

In this study, we explored the dynamics and the physiological role of LDs in neurons. Using *C. elegans* as a model, we found that ATGL-1/LID-1-mediated lipolysis autonomously regulates neuronal LD dynamics. ATGL-1 is the *C. elegans* homolog of mammalian ATGL, the rate-limiting enzyme of TAG hydrolysis, and LID-1 is the *C. elegans* homolog of mammalian CGI-58 (also named as ABHD5), which is the best known co-activator of ATGL (Lass, Zimmermann et al., 2006). Mutations in either CGI-58 or ATGL lead to neutral lipid storage disease (NLSD) with neurological abnormalities in human (Massa, Pozzessere et al., 2016, Schweiger, Lass et al., 2009). Importantly, defective neuronal lipolysis reduces PUFA-mediated touch sensation and protects neurons from hyperactivation-triggered neurodegeneration. The neuronal protective effect of *atgl-1* mutants is significantly enhanced by reduction of *de novo* PUFA synthesis and/or incorporation of PUFAs into phospholipids. Together, our results show that neuronal LDs participates in PUFA-mediated neural functions and neurodegeneration.

## Results

### LDs accumulate in neurons of both *atgl-1* and *lid-1* mutants

To observe LDs in worm neurons, we generated a neuron-specific GFP reporter line *xdIs109[Punc-119::PLIN1::GFP/rol-6]* expressing the LD surface protein PLIN1. PLIN1::GFP forms ring-like structures when expressed in tissues with LDs (Bi, Xiang et al., 2012, Liu, Li et al., 2014). In *xdIs109* animals, we mainly focused on the head region, which is the location of most neuronal soma. We found that there are very few GFP rings in *xdIs109* young adults (Fig. 1A). This indicates that similar to mammals, there are few LDs in *C. elegans* neurons under normal conditions. It has been reported that both aging and general obesity may increase LD accumulation in non-adipose tissues (Palikaras, Mari et al., 2017, Shimabukuro, Langhi et al., 2016, Zhou, Grayburn et al., 2000). To explore the influence of aging and overall fat content increase on neuronal LDs, we examined the *xdIs109* GFP pattern in 8-day-old wild-type adults, *daf-2(e1370)* mutants and *glp-1(e2141)* mutants. It is well known that the latter two mutants have an overall increase in lipid storage (O’Rourke, Soukas et al., 2009). We found that compared to young *xdIs109* controls, there is no significant increase of LD number in 8-day-old *xdIs109, daf-2(e1370); xdIs109* and *glp-1(e2141); xdIs109* (Fig. 1B). This indicates that aging and general obesity do not necessarily result in neuronal LD accumulation.

**Fig. 1.**
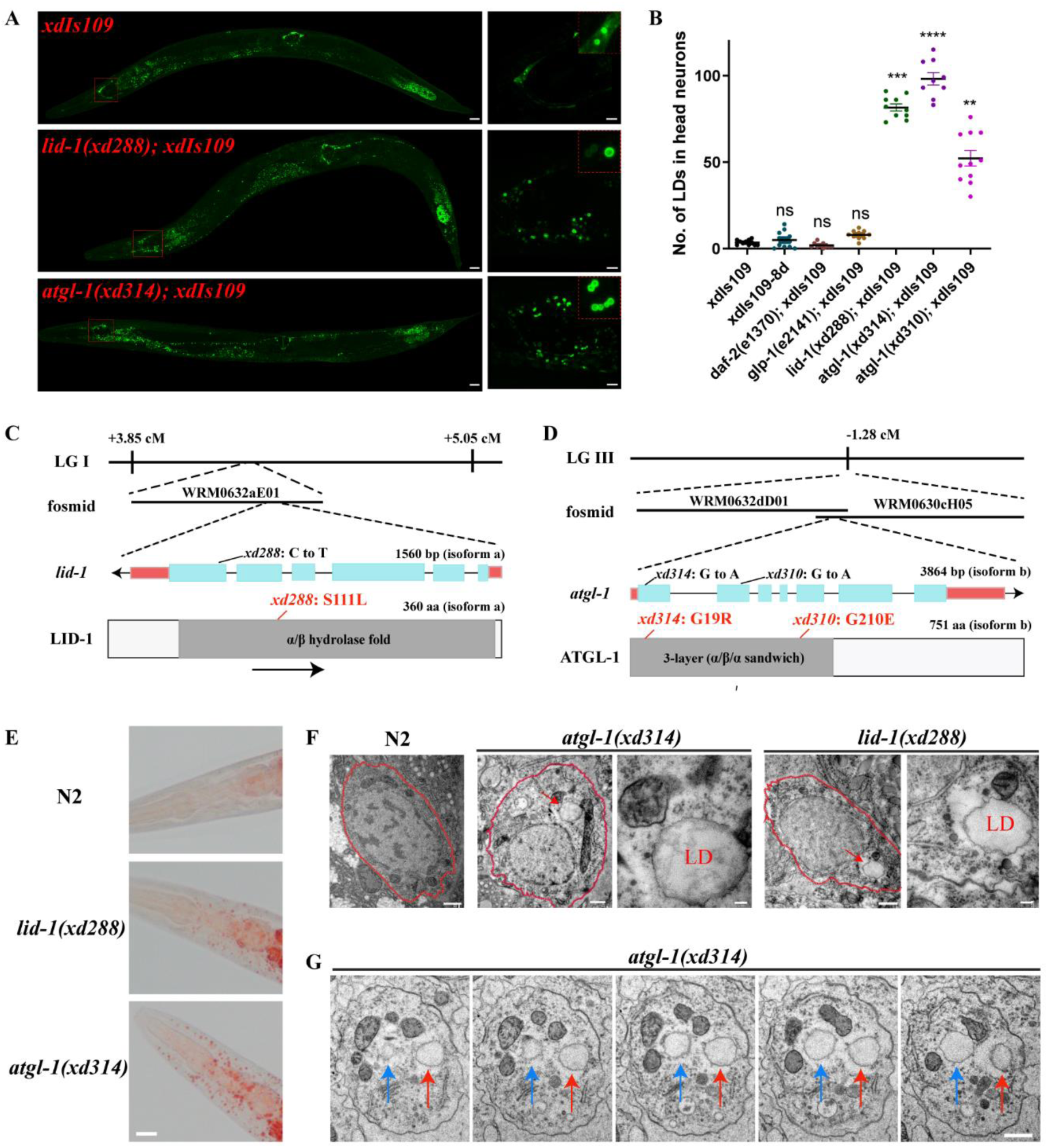
*lid-1(xd288)* and *atgl-1(xd314)* display ectopic LD accumulation in neurons. (A) *lid-1(xd288)* and *atgl-1(xd314)* mutants show LD accumulation in most neurons in *C. elegans*, including neurons in head region, tail and ventral nerve cord. *xdIs109* is a stable transgenic line with pan-neuronal expression of LD marker PLIN1::GFP. Scale bar: 20 μm. The right panels depict the head region. Scale bar: 5 μm. The insets in the right panels are enlarged views of PLIN1::GFP rings, representing LDs. (B) Quantification of the numbers of LDs in head region neurons (each dot represents one worm). The data was analyzed using one-way ANOVA with Dunnett’s multiple comparison test. * denotes the significant difference as compared to the control *xdIs109*. Error bars, SEM. n ≥ 9. (C-D) The genetic locus of *lid-1* and *atgl-1*. The coding regions are in light blue boxes and the noncoding regions are shown as lines. The UTRs are in red boxes. *xd288* is a point mutant of *lid-1* and causes a missense S111L mutation in the α/β hydrolase domain. *xd314* and *xd310* are point mutants of *atgl-1. xd314* harbors a missense G19R mutation and *xd310* harbors a missense G210E mutation. (E) Oil red O staining show dramatically increased neutral lipids in head region of *lid-1(xd288)* and *atgl-1(xd314)*. Scale bar: 10 μm. (F) EM images of head neurons. The red dash line marks outline of neurons. The red arrows mark LDs. Scale bar: 0.5 μm. The scale bar in enlarged images is 100 nm. (G) Serial sections of *atgl-1(xd314)* show LDs in neuron cell body. Scale bar: 0.5 μm.

To reveal the underlying mechanism(s) that control LD dynamics in neurons, we performed an EMS screen using the *xdIs109* marker to search for mutants with neuronal LD accumulation. We isolated the *xd288, xd310* and *xd314* mutants, which show LDs in many neurons in the head region, ventral nerve cord and tail (Fig. 1A). We also quantified LDs in the head region, and found that the number of LDs is increased significantly in these mutants (Fig. 1B).

Complementation tests divided these three mutants into two complementation groups. *xd288* complements both *xd310* and *xd314*, while *xd310* fails to complement *xd314*. Through SNP mapping, we narrowed down the region of the *xd288* mutation to chromosome I between +3.85 cM and +5.05 cM. Through fosmid transgenic rescue assays, we found that the WRM0632aE01 fosmid fully rescues the neuronal LD phenotype of *xd288*. The gene *lid-1* (lipid droplet protein 1), which encodes one of the *C. elegans* homologs of mammalian CGI-58, is found in this fosmid (Lee, Kong et al., 2014). Importantly, we found a C332T point mutation in *lid-1*, and this mutation leads to a missense S111L mutation of LID-1 (Fig. 1C). S111 is located in the putative α/β hydrolase domain of LID-1 and is conserved in *C. elegans, Drosophila*, mouse and human (Fig. S1A). Interestingly, S111 of LID-1 corresponds to S115 in human, which is mutated in NLSD (Ben Selma, Yilmaz et al., 2007).

A previous report showed that LID-1 binds to the *C. elegans* ATGL homolog, ATGL-1, and promotes ATGL-1-dependent lipolysis during fasting conditions (Lee et al., 2014). As expected, we found that *xd310* and *xd314* are mutants of *C. elegans atgl-1*. The *xd310* mutation causes a missense G210E change, while *xd314* causes a missense G19R change in ATGL-1 (Fig. 1D). G19 of ATGL-1 is conserved in *C. elegans, Drosophila*, mouse and human (Fig. S1B). The G19R and G210E mutations are located in an active (α/β/α) sandwich domain of ATGL-1, which is responsible for its enzymatic activity. Oil red O staining shows that both *lid-1(xd288)* and *atgl-1(xd314)* have more fat than controls, especially in the head region (Fig. 1E) and intestine (Fig. S1C). This suggests that, just like in NLSD, the overall fat content is increased in *lid-1* and *atgl-1* mutants. The recessive nature of the *xd288, xd310* and *xd314* mutations and the lipolytic function of ATGL and CGI-58 suggest that defective lipolysis leads to LD accumulation in neurons.

These mutants were isolated based on the appearance of neuronal LDs in the *xdIs109* background. To avoid potential interference from the overexpression of PLIN1::GFP, we used electron microscopy (EM) to observe neurons in *lid-1(xd288)* and *atgl-1(xd314)* animals without the *xdIs109* marker. LDs were easily found in the neuronal soma of *xd288* and *xd314* mutants, but not in wild type (Fig. 1F). We could even trace whole LDs in mutants from continuous EM serial sections (Fig. 1G). This demonstrates that *atgl-1(xd314)* and *lid-1(xd288)* mutants indeed have ectopic LDs in neurons.

### *atgl-1* autonomously regulates neuronal LDs

ATGL-1 is expressed strongly in worm intestine (Lee et al., 2014), but its expression in neurons has not been reported in detail. To observe the expression pattern of ATGL-1, we first mapped the promotor of *atgl-1*. The excess neuronal LD phenotype of *atgl-1(xd314)* was fully rescued by several overlapping *atgl-1(+)* fosmids. The overlapping region of the rescuing fosmids highlights a 3.4 Kb promoter region which may be important for the rescuing activity. We further found that an *atgl-1* genomic fragment with a 3 Kb promoter fully rescues the neuronal LD phenotype of *atgl-1(xd314)* (Fig. S2A and B). Using this 3 Kb promoter region to drive a GFP reporter, we examined the expression pattern of *atgl-1* in detail. The GFP fluorescence is bright in the intestine, consistent with a previous report (Lee et al., 2014). We also found weak expression in neurons in the nerve ring, ventral nerve cord and tail (Fig. 2A).

**Fig. 2.**
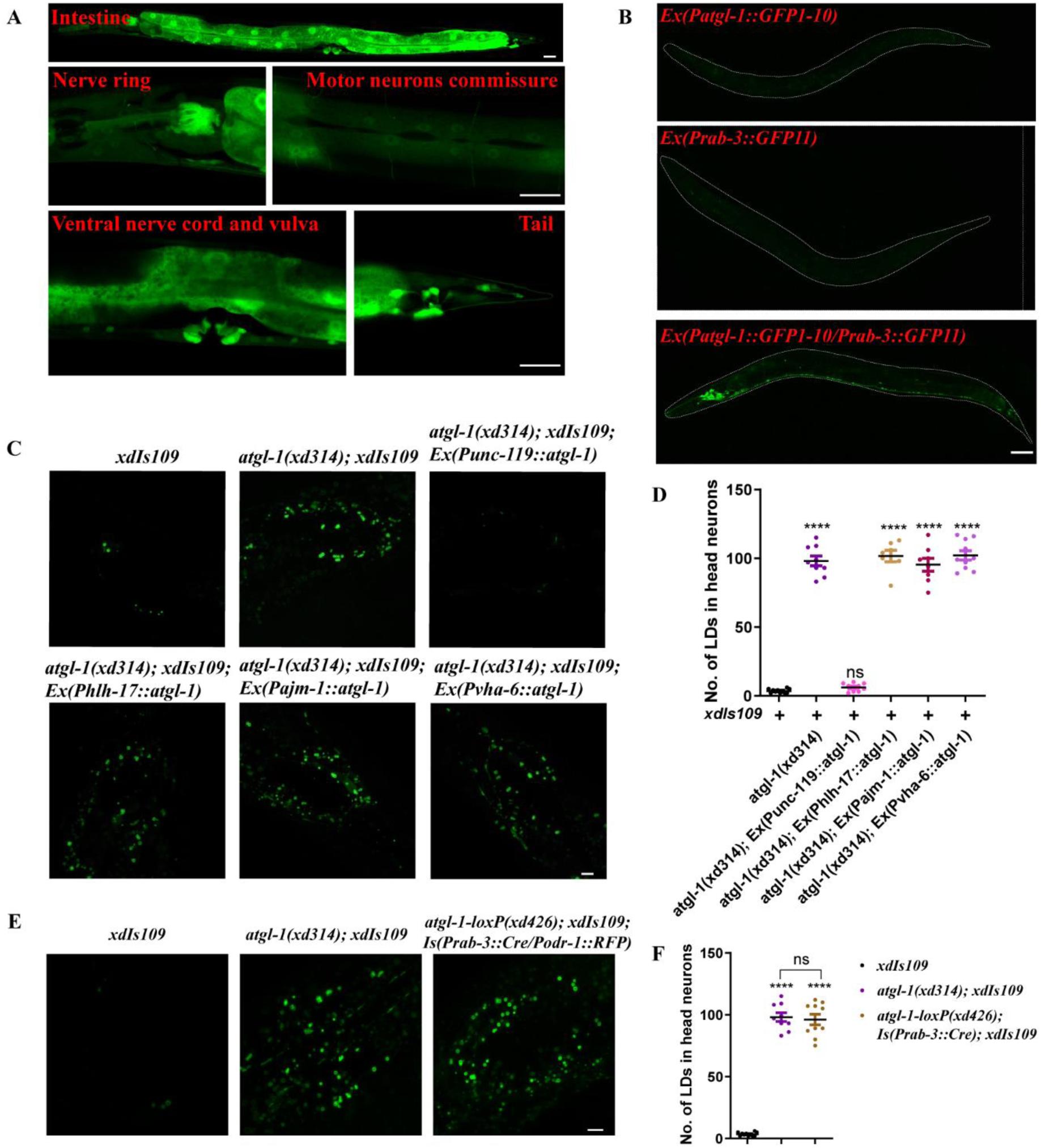
*atgl-1* autonomously regulates neuronal LD dynamics. (A) The expression pattern of *atgl-1* viewed by *Patgl-1::GFP* transcriptional fusion reporter. *Patgl-1::GFP* expression is high in intestine. Relatively low fluorescent signals are found in nerve ring, motor neuron commissure, ventral nerve cord and tail. Scale bar: 20 μm. (B) *atgl-1* is expressed in neurons. There is no fluorescent signal when expressing *Patgl-1::GFP1-10* and *Prab-3::GFP11* separately. With the co-expression of *Patgl-1::GFP1-10* and *Prab-3::GFP11*, the GFP signals are found in head region, tail and ventral nerve cord, where most neuronal cell bodies are located. The white dash lines mark the outline of worms. Scale bar: 50 μm. (C) Visualizing LDs in head neurons with *xdIs109* reporter in different genetic backgrounds. The accumulation of LDs in head neurons of *atgl-1(xd314)* mutants can be rescued by expressing wild-type *atgl-1* in neurons, but not in intestine, glia or hypodermis. Scale bar: 5 μm. (D) Quantification of the number of LDs in head neurons in different genetic backgrounds. Each dot represents one worm. The data was analyzed using one-way ANOVA with Dunnett’s multiple comparison test. * denotes the significant difference as compared to the control *xdIs109*. Error bars, SEM. n ≥ 7. ns: not statistically significant. (E) Neuron-specific knockout of *atgl-1* causes LD accumulation in neurons, similar to *atgl-1(xd314)* mutants. Scale bar: 5 μm. (F) Quantification of the number of LDs in head neurons in different genetic backgrounds. The data was analyzed using one-way ANOVA with Bonferroni’s multiple comparison test. * denotes the significant difference as compared to the control *xdIs109*. Error bars, SEM. n ≥ 9. ns: not statistically significant.

To specifically analyze ATGL-1 expression in neurons, we used the self-complementing split GFP system. Split GFP is composed of two separate GFP fragments (a 15-amino acid fragment, GFP11, and a 215-amino acid fragment, GFP1-10), which are expressed separately and are able to associate spontaneously to form fluorescent GFP (Cabantous, Terwilliger et al., 2005). As controls, transgenic animals expressing either *Patgl-1::GFP1-10* or pan-neuronal *Prab-3::GFP11* do not exhibit GFP fluorescence (Fig. 2B). The GFP fluorescence is clearly seen in neurons, including head region, ventral nerve cord and tail neurons, in animals expressing both *Patgl-1::GFP1-10* and pan-neuronal *Prab-3::GFP11* (Fig. 2B). This result provides further evidence for the neuronal expression of ATGL-1.

To investigate whether *atgl-1* functions autonomously or non-autonomously, we performed tissue-specific rescue and tissue-specific knockout experiments. We found that neuronal *Punc-119::atgl-1* expression, but not glial *Phlh-17::atgl-1*, hypodermal *Pajm-1::atgl-1* or intestinal *Pvha-6::atgl-1* expression, rescues the neuronal LD phenotype of *atgl-1(xd314); xdIs109* (Fig. 2C and D). We also generated a neuron-specific knockout of *atgl-1* using the Cre-loxP system. We first used CRISPR-Cas9 to insert two loxP sites flanking the *atgl-1* coding region, and then we expressed Cre in neurons to specifically knock out *atgl-1* in neurons (Fig. S2C). We found that similar to *atgl-1(xd314*) mutants, neuronal-specific deletion of *atgl-1 [atgl-1-loxP(xd426); xdIs182 (Prab-3::Cre/Podr-1::RFP)]* causes LD accumulation in neurons (Fig. 2E and F). Together, these results demonstrate that ATGL-1 acts autonomously in neurons to prevent neuronal LD accumulation.

### Paralogs of ATGL-1 and LID-1 do not affect neuronal LD dynamics

The above results indicate that ATGL-1/LID-1-mediated lipolysis prevents the appearance of visible LDs in neurons. In contrast to lipolysis, lipogenesis promotes lipid storage. We wondered whether overexpression of lipogenesis-related genes in neurons could cause LD accumulation. DGAT1 and DGAT2 are key enzymes in the synthesis of TAG from diacylglycerol (DAG) (Fig. 3A). DGAT1 has only one homolog, MBOA-2, in *C. elegans*. DGAT2 has four homologs (DGAT-2/F59A1.10, K07B1.4, DGTR-1/W01A11.2 and Y53G8B.2) in *C. elegans* (Fig. 3B). We pan-neuronally overexpressed *mboa-2* or *K07B1.4*. In these transgenic animals, there are ectopic LDs in neurons as revealed by the *xdIs109* reporter (Fig. 3C). Therefore, both elevating lipogenesis or reducing lipolysis leads to LD accumulation in neurons. Together, these results indicate that neurons have the ability to form and hydrolyze LDs. The lack of visible LDs in neurons reflects the dominance of lipolysis over lipogenesis under normal conditions.

**Fig. 3.**
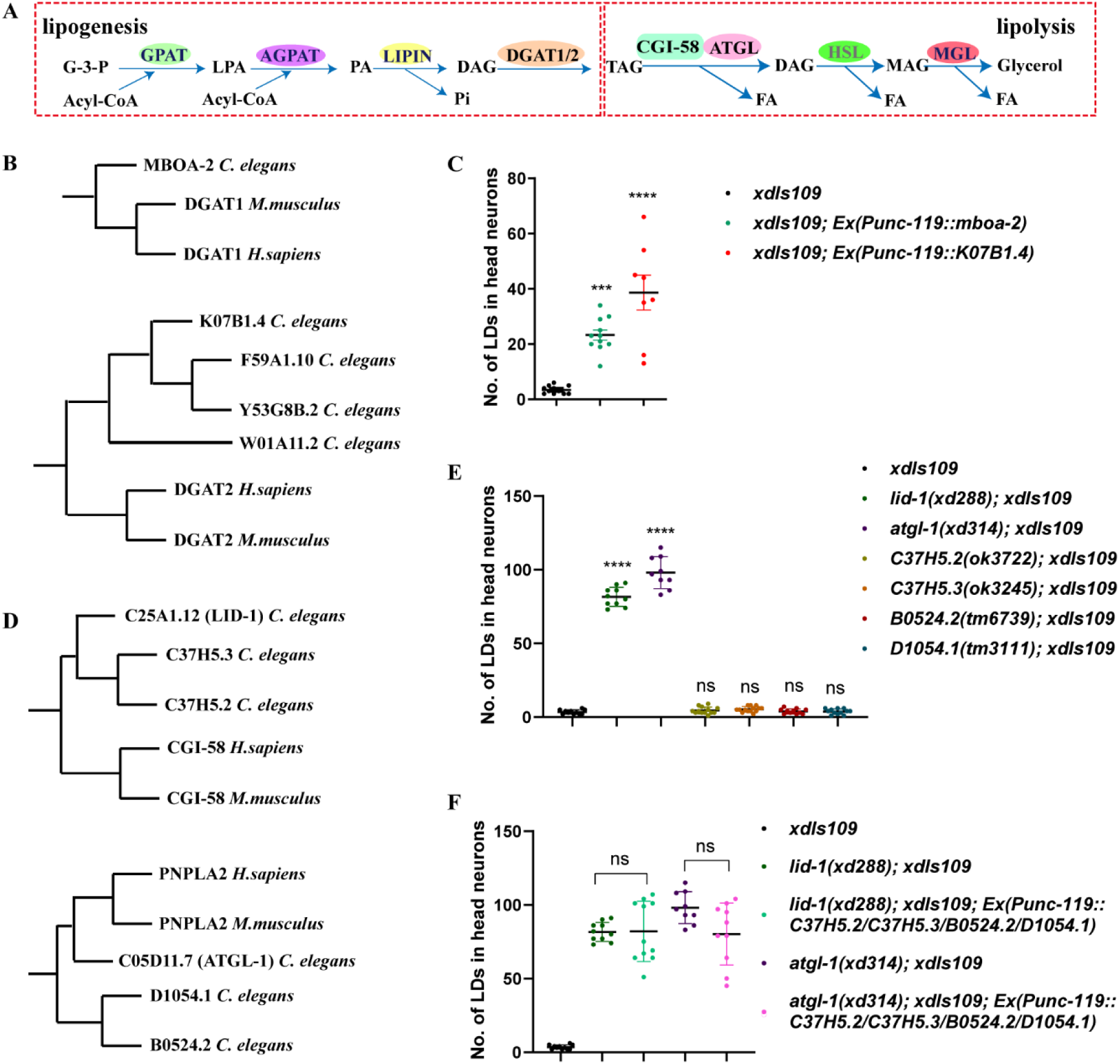
LID-1/ATGL-1 mediated lipolysis and DAGT1/2 mediated lipogenesis regulate LD dynamics in neurons. (A) The pathway of lipolysis and lipogenesis. G-3-P: glycerol-3-phosphate. LPA: lysophosphatidic acid. PA: phosphatidic acid. DAG: diacylglycerol. TAG: triacylglycerol. MAG: monoacylglycerol. FA: fatty acid. (B) The evolutionary phylogenetic tree of DGAT1 and DGAT2. (C) Overexpressing the homologs of DGAT1/2 in neurons of *C. elegans* leads to LD accumulation in neurons. Each dot represents one worm. The data was analyzed using one-way ANOVA with Dunnett’s multiple comparison test. * denotes the significant difference as compared to the control *xdIs109*. Error bars, SEM. n ≥ 8. (D) The evolutionary phylogenetic tree of LID-1 and ATGL-1. (E) Mutants of LID-1 or ATGL-1 paralogs do not affect LD dynamics in neurons. Each dot represents one worm. The data was analyzed using one-way ANOVA with Dunnett’s multiple comparison test. * denotes the significant difference as compared to the control *xdIs109*. Error bars, SEM. n ≥ 9. ns: not statistically significant. (F) Neuronal expression of LID-1 and ATGL-1 paralogs do not rescue *lid-1(xd288)* or *atgl-1(xd314)*. Each dot represents one worm. The data was analyzed using one-way ANOVA with Tamhane’s T2 multiple comparison test. * signifies the significant difference between the groups under the crossbar. Error bars, SEM. n ≥ 9. ns: not statistically significant.

There are two LID-1 paralogs (C37H5.2 and C37H5.3) and two ATGL-1 paralogs (B0524.2 and D1054.1) in *C. elegans* (Fig. 3D). Although LID-1 is reported to regulate lipolysis by interacting with ATGL-1 (Lee et al., 2014), *C37H5.3*, also named as *cgi-58*, had been shown to promote ATGL-1 activity at the dauer stage when the protein stability of ATGL-1 is negatively regulated via phosphorylation by AMPK (Narbonne & Roy, 2009, Xie & Roy, 2015a, Xie & Roy, 2015b). To test whether ATGL-1 and LID-1 paralogs affect LD homeostasis in neurons, we examined the neuronal LD phenotypes of *C37H5.2(ok3722), C37H5.3(ok3245), D1054.1(tm3111)* and *B0524.2(tm6739)* mutants with the *xdIs109* marker. Similar to wild type, there are almost no LDs in neurons in these mutants (Fig. 3E). These mutants are all deletion alleles (Fig. S3), and even though we are not sure whether they are null alleles, their phenotypes indicate that the ATGL-1/LID-1 pair plays the main role in regulating neuronal lipolysis and neuronal LD dynamics.

Next, we explored whether these paralogs can substitute for ATGL-1 or LID-1 in neurons. We overexpressed these paralogs in neurons specifically to examine whether they could rescue the neuronal LD phenotype of *atgl-1(xd314); xdIs109* or *lid-1(xd288); xdIs109*. Because these paralogs may function in pairs, like ATGL-1 and LID-1, we overexpressed all four paralogs together in neurons. We found that they could not rescue the neuronal LD phenotype of either *atgl-1(xd314); xdIs109* or *lid-1(xd288); xdIs109* (Fig. 3F). Together, these results indicate the importance and the specificity of the ATGL-1/LID-1 partnership in regulating neuronal LD dynamics.

### *atgl-1* and *lid-1* affect gentle touch sensation and genetically interact with the *de novo* PUFA biosynthesis pathway

The above results show that lipolysis restricts the appearance of visible LDs in neurons. We next examined the neuronal consequence of defective lipolysis. LDs are temporary storage places for excess lipids. Since lipids are important building blocks for membranes, we initially focused on examining whether *atgl-1* affects neuronal morphology. The PVD neuron has very complex neurites, especially dendrites, which may require lots of lipids for membrane biogenesis. We did not find any morphological difference between wild type and *atgl-1(xd314)* or *lid-1(xd288)* mutants based on observations with the PVD marker *wdIs52(P*_*F49H12.4*_::*GFP)* (Fig. S4). This suggested that the appearance of LDs may not affect the gross morphology of neurons.

A previous report showed that when PUFAs are diverted away from membranes to the core of LDs, the expansion of LDs inhibits the oxidation of PUFAs and reduces the toxic effect of PUFAs (Bailey et al., 2015). Therefore, the presence of LDs in neurons may limit the availability of PUFAs and affect PUFA-associated neuronal events. PUFAs affect the gentle touch sensation and mechanoelectrical transduction in *C. elegans* touch receptor neurons (Kahn-Kirby, Dantzker et al., 2004, Vasquez, Krieg et al., 2014). There are ectopic LDs in touch neurons in *atgl-1(xd314)* as revealed by stimulated Raman scattering (SRS) microscopy (Fig. 4A). To examine whether the gentle touch sensation is affected in *atgl-1(xd314)* and *lid-1(xd288)* mutants, we measured the touch sensitivity by a ten-trial touch assay (Hart, 2006). *mec-4(u253)* and N2 were used as the positive control and negative control, respectively. Compared to the strong touch sensation defect in *mec-4(u253)* mutants, touch sensation is partially impaired in *atgl-1(xd314)* and *lid-1(xd288)* mutants. Importantly, expression of neuronal specific *P*_*unc-119*_::*atgl-1* and *P*_*unc-119*_::*lid-1* fully rescued the touch sensation defect in *atgl-1(xd314)* and *lid-1(xd288)*, respectively (Fig.4B). This suggests that neuronal lipolysis is required for normal gentle touch sensation.

**Fig. 4.**
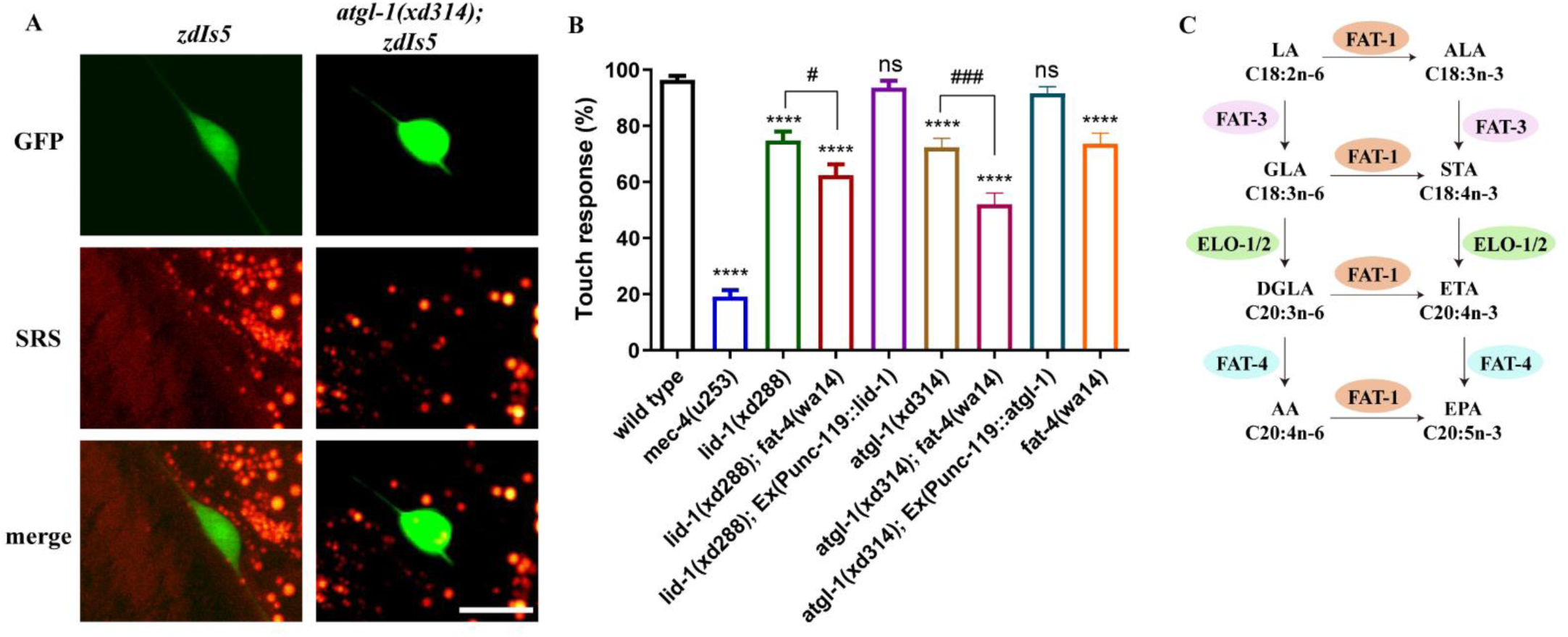
*atgl-1(xd314)* and *lid-1(xd288)* show gentle touch sensation defect. (A) LDs accumulate in touch neurons in *atgl-1(xd314)* visualized by SRS (Stimulated Raman scattering). The *zdIs5(P*_*mec-4*_:: *GFP)* reporter marks touch neurons. This shows that there is LD accumulation in touch neurons in *atgl-1(xd314)*. Scale bar: 10 μm. (B) *atgl-1(xd314)* and *lid-1(xd288)* show gentle touch sensation defect, which could be rescued by the neuron-specifically expression of *atgl-1* and *lid-1*, respectively. The mutation of *fat-4* enhances the touch sensation defect in *atgl-1(xd314)* and *lid-1(xd288)*. The data was analyzed using one-way ANOVA with Turkey’s multiple comparison test. * denotes the significant difference as compared to the control N2. “NS” and “#” signify the significant difference between the groups under the crossbar. Error bars, SEM. n =25. (C) The pathway of PUFA synthesis in *C. elegans*. LA: linoleic acid. ALA: alpha linolenic acid. GLA: gamma linoleic acid. STA: stearidonic acid. DGLA: dihommo gamma linoleic acid. ETA: eicosatetraenoic acid. AA: arachidonic acid. EPA: eicosapentaenoic acid. Fatty acids are also indicated by chemical abbreviations. For example, the abbreviation C18:2n-6 means 18 carbons with two double bonds and the first double bond is located at ω-6.

In the *de novo* PUFA biosynthetic pathway, *fat-4* is responsible for the synthesis of arachidonic acid (AA, C20:4 (20 carbons with four double bonds)) and eicosapentaenoic acid (EPA, C20:5) (Fig.4C) (Watts & Browse, 2002). Mutation of *fat-4* leads to mild gentle touch sensation defects similar to those in *atgl-1(xd314)* and *lid-1(xd288)* mutants (Fig.4B) (Vasquez et al., 2014). We then investigated the genetic relationship between ATGL-1/LID-1-mediated lipolysis and *de novo* synthesis of PUFAs by double mutant analysis. Interestingly, the *fat-4(wa14)* mutation enhances the touch sensation defect of both *atgl-1(xd314)* and *lid-1(xd288)* single mutants (Fig. 4B), which suggests that ATGL/LID-1-regulated neuronal lipolysis participates in PUFA-mediated touch sensation.

### *atgl-1(xd314)* and *lid-1(xd288)* mutants have reduced neuron hyperactivation-triggered neurodegeneration

Glial LDs are reported to be involved in neurodegeneration in *Drosophila* (Liu et al., 2017, Liu et al., 2015). In particular, to avoid fatty acid toxicity in hyperactivated neurons, fatty acids are transported from neurons into glia and stored in LDs before detoxification through mitochondrial oxidation (Ioannou et al., 2019). These findings prompted us to investigate whether the appearance of neuronal LDs in *atgl-1* and *lid-1* mutants alleviates neurodegeneration. *mec-4* encodes the ion channel protein MEC-4 and dominant negative mutants of *mec-4*, often referred to as *mec-4(d)*, have been widely used as models of neuron hyperactivation-triggered neurodegeneration (Calixto, Jara et al., 2012, Driscoll & Chalfie, 1991). The neurodegeneration in *mec-4(d)* occurs as early as the embryonic stage, and by the L4 stage the majority (63 ± 2%) of *mec-4(d)* animals only have two touch neurons left as viewed by the *zdIs5* marker, which labels six touch sensory neurons in wild type (Fig. 5A). We found that *atgl-1* mutation reduces the neuronal degeneration of *mec-4(d)* (Fig. 5A and B). Specifically, the proportion of animals with three or more surviving touch neurons increased from 6 ± 1% in *mec-4(d)* to 28 ± 1% in *atgl-1(xd314)*; *mec-4(d)* worms (Fig. 5C). Similarly, *lid-1* mutation also reduced the neuronal loss of *mec-4(d)*. The proportion of animals with three or more surviving touch neurons increased to 23 ± 2% in *lid-1(xd288)*; *mec-4(d)* worms (Fig. 5C). Moreover, the reduction of *mec-4(d)-*triggered neurodegeneration by *atgl-1* and *lid-1* mutations can be reversed by neuronal specific expression of wild-type *atgl-1* and *lid-1*, respectively (Fig. 5C). These data indicate that defective neuronal lipolysis reduces neuron hyperactivation-triggered neurodegeneration.

**Fig. 5.**
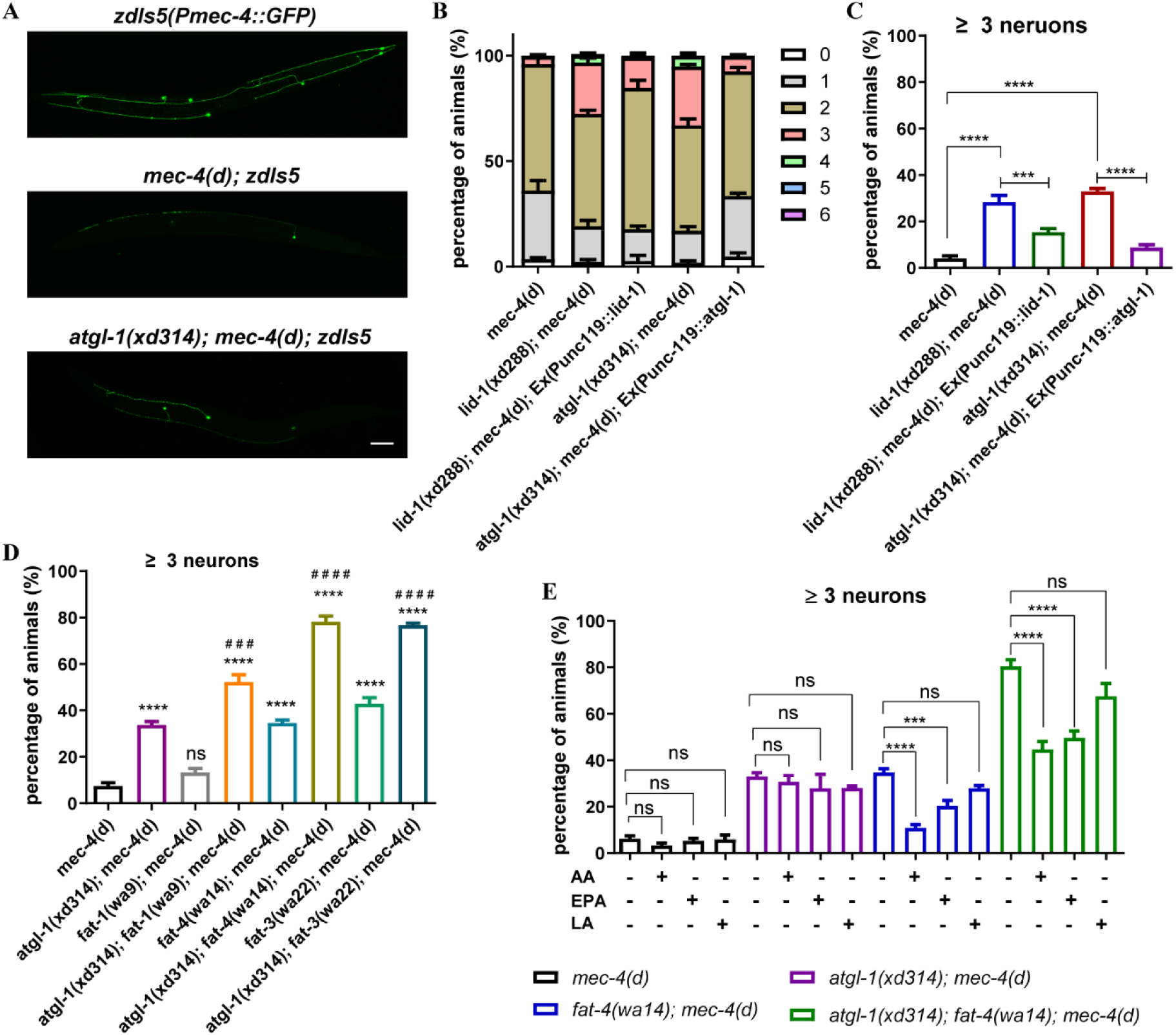
PUFA synthesis defect significantly enhances the beneficial effect of *atgl-1* mutants in preventing the neurodegeneration of *mec-4(d)*. (A) Image of touch neurons viewed by *zdIs5(Pmec-4::GFP)* in different genetic backgrounds. Scale bar: 20 μm. (B) Quantification of the percentage of animals with 0, 1, 2, 3, 4, 5 and 6 surviving touch neurons in different genetic backgrounds. At least 50 animals per strain were analyzed in each experiment. Number of experiments was 5. (C) The percentage of worms that have three or more surviving touch neurons in different genetic backgrounds. *lid-1(xd288)* and *atgl-1(xd314)* have significantly increased the number of surviving neurons. Neuron-specifically expression of *atgl-1* or *lid-1* reverses the protecting effect of *atgl-1* or *lid-1* mutation. The data was analyzed using one-way ANOVA with Turkey’s multiple comparison test. Error bars, SEM. ns: not statistically significant. Number of experiments n = 5, with at least 50 animals per strain analyzed in each experiment. (D) The percentage of worms that have three or more surviving touch neurons in different genetic backgrounds. Mutants of *fat-1, fat-3* and *fat-4* significantly enhance the suppression effect of *atgl-1(xd314)* in preventing the neurodegeneration of *mec-4(d)*. The data was analyzed using one-way ANOVA with Bonferroni’s multiple comparison test. # denotes the significant difference as compared to *atgl-1(xd314); mec-4(d)*. * signifies the significant difference as compared to *mec-4(d)*. Error bars, SEM. Number of experiments n ≥ 3, with at least 50 animals per strain analyzed in each experiment. (E) The percentage of worms that have three or more surviving touch neurons in different genetic backgrounds supplemented with different fatty acids. AA and EPA but not LA significantly enhance the neurodegeneration triggered by *mec-4(d)*. The data was analyzed using one-way ANOVA with Dunnett’s multiple comparison test. “ns” and “*” signify the significant difference between the groups under the crossbar. Error bars, SEM. Number of experiments n ≥ 3, with at least 50 animals per strain analyzed in each experiment.

### Defects in neuronal lipolysis and *de novo* PUFA biosynthesis synergistically decrease the neuronal loss caused by *mec-4(d)*

Hyperactivated neurons are vulnerable because the peroxidation of fatty acids, in particular PUFAs, can result in neurodegeneration (Ioannou et al., 2019). *fat-1, fat-3* and *fat-4* encode key enzymes for *de novo* synthesis of different PUFAs in *C. elegans* (Watts & Browse, 2002, Watts & Ristow, 2017). Consistent with the potential involvement of PUFAs in neuron hyperactivation-triggered neurodegeneration, we found that *mec-4(d)-*triggered neurodegeneration was significantly reduced in *fat-4(wa14)* and *fat-3(wa22)* mutants but not *fat-1(wa9)* (Fig. 5D). The proportion of animals that with three or more surviving touch neurons increased from 6 ± 1% in *mec-4(d)* to 35 ± 1% in *fat-4(wa14)*; *mec-4(d)* and 43 ± 3% in *fat-3(wa22)*; *mec-4(d)* worms. These results indicate that reducing the biosynthesis of PUFAs alleviates the neuronal loss of *mec-4(d)*.

Since ATGL-1/LID-1-mediated neuronal lipolysis is involved in PUFA-mediated touch sensation (Fig. 5A and B), and both *atgl-1* and *lid-1* mutants have reduced *mec-4(d)*-induced neurodegeneration (Fig. 4B), we next explored the neuroprotective effect of double mutations in neuronal lipolysis and *de novo* synthesis of PUFAs. Interestingly, *atgl-1(xd314); fat-1(wa9), atgl-1(xd314); fat-3(wa22)* and *atgl-1(xd314); fat-4(wa14)* double mutants all have significantly reduced *mec-4(d)-*triggered neurodegeneration compared to single mutants (Fig. 5D). The proportion of animals with three or more surviving touch neurons increased from 6 ± 1% in *mec-4(d)* to 52 ± 3% in *fat-1(wa9)*; *atgl-1(xd314); mec-4(d)*, 78 ± 2% in *fat-4(wa14)*; *atgl-1(xd314); mec-4(d)* and 77 ± 1% in *fat-3(wa22)*; *atgl-1(xd314); mec-4(d)* worms. Therefore, blocking *de novo* PUFA synthesis and neuronal lipolysis together greatly protects neurons from degeneration caused by *mec-4(d)*.

### The PUFAs AA and EPA promote the neurodegeneration of *mec-4(d)*

Since *fat-3(wa22)* and *fat-4(wa14)* show similar neural protection phenotypes (Fig 5D), we reasoned that PUFAs, such as AA (C20:4, n-6) and EPA (C20:5, n-3) or their derivatives, may promote *mec-4(d)-*induced neurodegeneration. We fed worms on NGM medium supplemented with different fatty acids. *fat-4(wa14); mec-4(d)* and *fat-4(wa14); atgl-1(xd314); mec-4(d)* animals show a significant increase of neurodegeneration when grown in AA-supplemented NGM compared with normal NGM (Fig. 5E). Feeding worms with EPA, but not linoleic acid (LA, C18:2), had a similar effect (Fig. 5E). In contrast, feeding worms with a mixture containing two saturated fatty acids (palmitic acid (PA, C16:0) and stearic acid (SA, C18:0)) and one monounsaturated fatty acid (oleic acid (OA, C18:1)) did not affect the neurodegeneration phenotype in *fat-4(wa14); mec-4(d)* and *fat-4(wa14); atgl-1(xd314); mec-4(d)* (Fig. S5). These results indicate that the PUFAs AA and EPA promote the neurodegeneration of *mec-4(d)*.

### PUFAs promote neurodegeneration through incorporation into phospholipids

Elevated ROS is proposed to be a key event for neurodegeneration triggered by neuron hyperactivation in both *C. elegans* and mammals (Calixto et al., 2012, Ioannou et al., 2019, Sangaletti, D’Amico et al., 2017). We then tested whether ROS contributes to PUFA-mediated neurodegeneration. We found that *sod-2* mutation did not significantly decrease the number of surviving neurons in *atgl-1(xd314); mec-4(d), fat-4(wa14); mec-4(d)*, or *fat-4(wa14); atgl-1(xd314); mec-4(d)* (Fig. S6A). Furthermore, overexpression of the ROS scavenger SOD-1, SOD-2, SOD-3 or SOD-4 in touch neurons did not affect *mec-4(d)*-triggered neurodegeneration (Fig. S6B). Moreover, when we fed *mec-4(d)* mutants with the antioxidant N-acetylcysteine (NAC), the survival of neurons was still not significantly increased (Fig. S6C). In addition, there was no change in the number of surviving neurons in *atgl-1(xd314)*; *mec-4(d), fat-4(wa14); mec-4(d)* and *atgl-1(xd314); fat-4(wa14); mec-4(d)* following treatment with N-acetylcysteine (NAC) (Fig. S6C). Therefore, ROS may not play a major role in neuronal lipolysis and PUFA-mediated neuronal loss caused by *mec-4(d)*.

To further investigate how PUFAs promote neurodegeneration, we performed a lipidomic analysis of *mec-4(d), atgl-1(xd314); mec-4(d), fat-4(wa14); mec-4(d)* and *fat-4(wa14); atgl-1(xd314); mec-4(d)* to find correlations between the level of PUFA-containing lipids and the severity of neurodegeneration. In *fat-4* mutants, C20:3 or C20:4-containing lipids are increased significantly while C20:5-containing lipids are decreased dramatically when normalized to total polar lipids (Fig. S7). This is consistent with the role of FAT-4 in generating EPA. Furthermore, since our analysis cannot distinguish between eicosatetraenoic acid (ETA, C20:4 n-3) and AA (C20:4 n-6), it is plausible that the increased C20:4 in *fat-4* mutants is ETA, as revealed in a previous study (Watts & Browse, 2002). Moreover, despite the increased level of C20:4-containing lipids, the percentage of C20:4-containing phospholipids is reduced in *fat-4* mutants, while the percentage of C20:4-containing TAG is increased significantly (Fig. 6A&B). Therefore, *fat-4* mutation reduces the levels of some PUFAs and may also alter the partitioning of PUFAs between phospholipids and TAG.

**Fig. 6.**
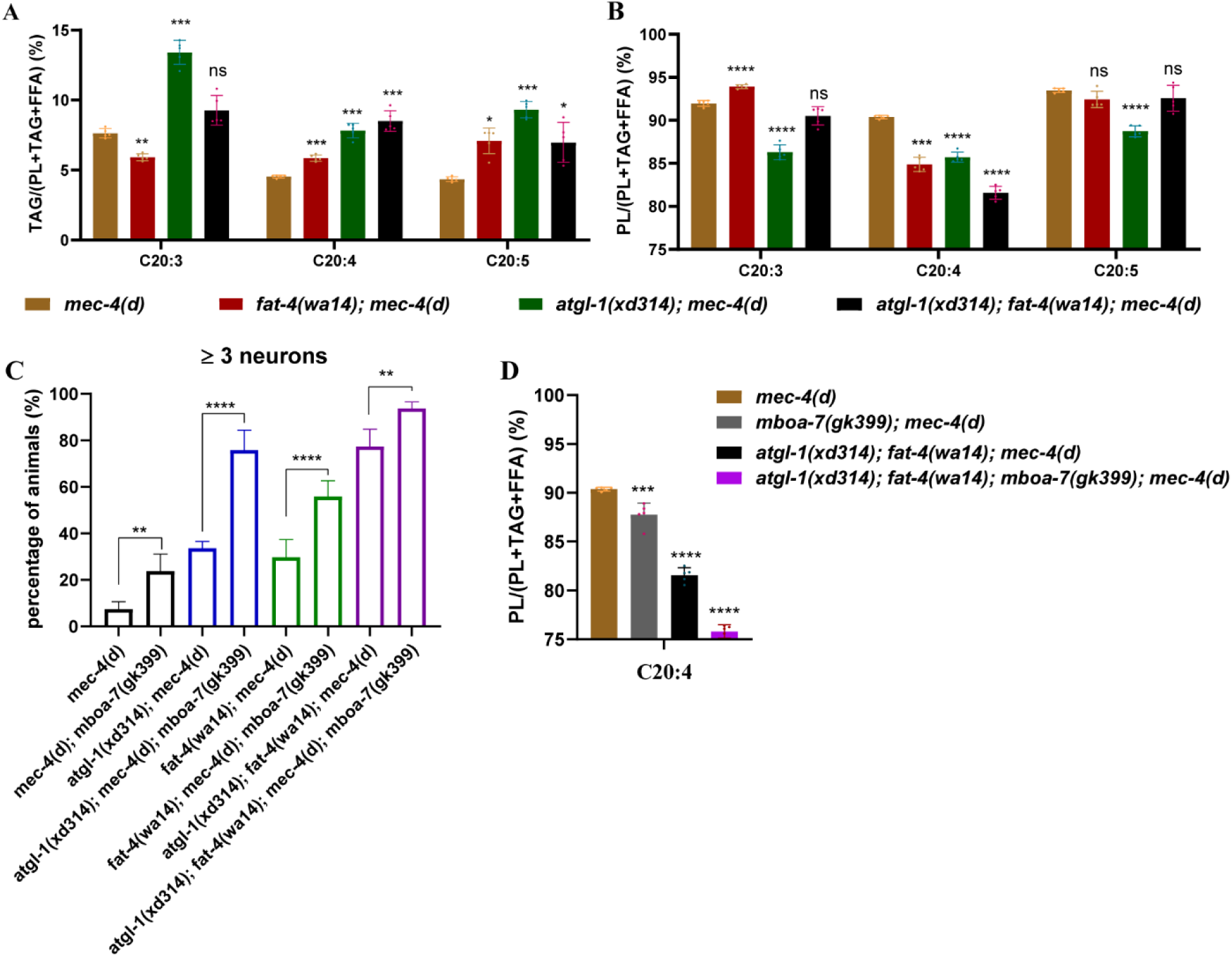
PUFA containing phospholipids regulate neurodegeneration of *mec-4(d)*. (A) The percentage of C20:3-, C20:4- or C20:5-containing TAGs to total lipids containing according PUFA in different genetic backgrounds. The percentage of C20:4-containing TAGs to total lipids containing C20:4 is significantly increased in *atgl-1* and *fat-4* single mutants. The percentage is further increased in *atgl-1* and *fat-4* double mutant. (B) The percentage of C20:3-, C20:4- or C20:5-containing phospholipids to total lipids containing according PUFA in different genetic backgrounds. The percentage of C20:4-containing phospholipids to total lipids containing C20:4 is significantly decreased in *atgl-1* and *fat-4* single mutants. The percentage is further decreased in *atgl-1* and *fat-4* double mutant. (A-B) FFA: free fatty acids; TAG: triacylglycerol; PL: phospholipids. The data was analyzed using two-way ANOVA with Dunnett’s multiple comparison test. * signifies the significant difference as compared to *mec-4(d)*. Error bars, SEM. n=5. ns: not statistically significant. (C) The percentage of worms that have three or more surviving touch neurons in different genetic backgrounds. *mboa-7* mutation significantly increases the percentage of three and more surviving touch neurons in *mec-4(d), atgl-1(xd314); mec-4(d), fat-4(wa14); mec-4(d)* and *atgl-1(xd314); fat-4(wa14); mec-4(d)*. The data was analyzed using one-way ANOVA with Bonferroni’s multiple comparison test. “*” signifies the significant difference between the groups under the crossbar. Error bars, SEM. n ≥ 4. ns: not statistically significant. (D) The percentage of C20:4-containing phospholipids to total lipids that have C20:4 is dramatically decreased in *mboa-7(gk399); atgl-1(xd314); fat-4(wa14); mec-4(d)*. The data was analyzed using one-way ANOVA with Dunnett’s multiple comparison test. * signifies the significant difference as compared to *mec-4(d)*. Error bars, SEM. n=5.

Compared to *fat-4* mutants, the levels of C20:3, C20:4, or C20:5-containing lipids are slightly increased in *atgl-1* mutants, when normalized to total polar lipids (Fig. S7). Interestingly, consistent with a storage role for LDs, the percentages of C20:3, C20:4, or C20:5-containing TAGs are significantly increased, while the percentages of C20:3, C20:4, or C20:5-containing phospholipids are significantly decreased in *atgl-1* mutants (Fig. 6A&B). Therefore, *atgl-1* mutation alters the partitioning of PUFAs into phospholipids and TAG.

In *atgl-1; fat-4* double mutants, there were similar changes of C20:3, C20:4 and C20:5 fatty acids as in *fat-4* single mutants. However, in the double mutants, we also noticed persistent and further changes in PUFA partitioning between phospholipids and TAG, in particular the partitioning of C20:4. There is a further reduction of the percentage of C20:4-containing phospholipids in *atgl-1; fat-4* double mutants compared to either single mutant (Fig. 6A&B). The altered partitioning of PUFAs between phospholipids and TAG correlates well with the moderate neuroprotective effects in *atgl-1* and *fat-4* single mutants and the strong neuroprotective effects in *atgl-1*; *fat-4* double mutants. These results suggest that PUFA-containing phospholipids may promote neurodegeneration.

We further explored the hypothesis that PUFA-containing phospholipids contribute to *mec-4(d)*-induced neurodegeneration. MBOA-7 incorporates PUFAs into phospholipids, especially PI (Lee, Inoue et al., 2008). We found that *mboa-7* mutation alone significantly increased the percentage of *mec-4(d)* animals with ≥3 surviving touch neurons (from ∼5% to ∼20%) (Fig. 6C). Moreover, *mboa-7* mutation further increases neuron survival in *atgl-1(xd314); mec-4(d), fat-4(wa14); mec-4(d)* and *atgl-1(xd314); fat-4(wa14); mec-4(d)* (Fig. 6C and Fig. S8). The enhancement of the neuroprotective effect is probably because the *mboa-7* mutation reduces PUFA incorporation into phospholipids. Consistent with that, the percentage of C20:4-containing phospholipids is further decreased in *atgl-1(xd314); fat-4(wa14); mboa-7(gk399); mec-4(d)* mutants compared with *atgl-1(xd314); fat-4(wa14); mec-4(d)* mutants (Fig. 6D). Together, these results indicate that PUFAs promote neuron hyperactivation-mediated neurodegeneration of *mec-4(d)* through their incorporation into phospholipids.

## Discussion

This study addresses two questions: why are there normally no LDs in neurons, and what are the consequences of LD accumulation in neurons? We reveal that the balance of lipolysis and lipogenesis regulates neuronal LD dynamics. In particular, we show that defective ATGL-1/LID-1-mediated lipolysis causes LD accumulation autonomously in *C. elegans* neurons. Defective neuronal lipolysis affects normal gentle touch sensation, while importantly, it alleviates neurodegeneration triggered by neuron hyperactivation. The neuroprotective effect is synergistically enhanced when *de novo* biosynthesis of PUFA is blocked. Lastly, the incorporation of PUFAs into phospholipids likely underlies neuronal lipolysis and PUFA-mediated neurodegeneration. Therefore, both neuronal LD dynamics and *de novo* PUFA synthesis regulate the availability of PUFAs, which may be incorporated into phospholipids to maintain the proper function of neurons under normal conditions and to promote neurodegeneration under stress conditions (Fig. 7).

**Fig. 7.**
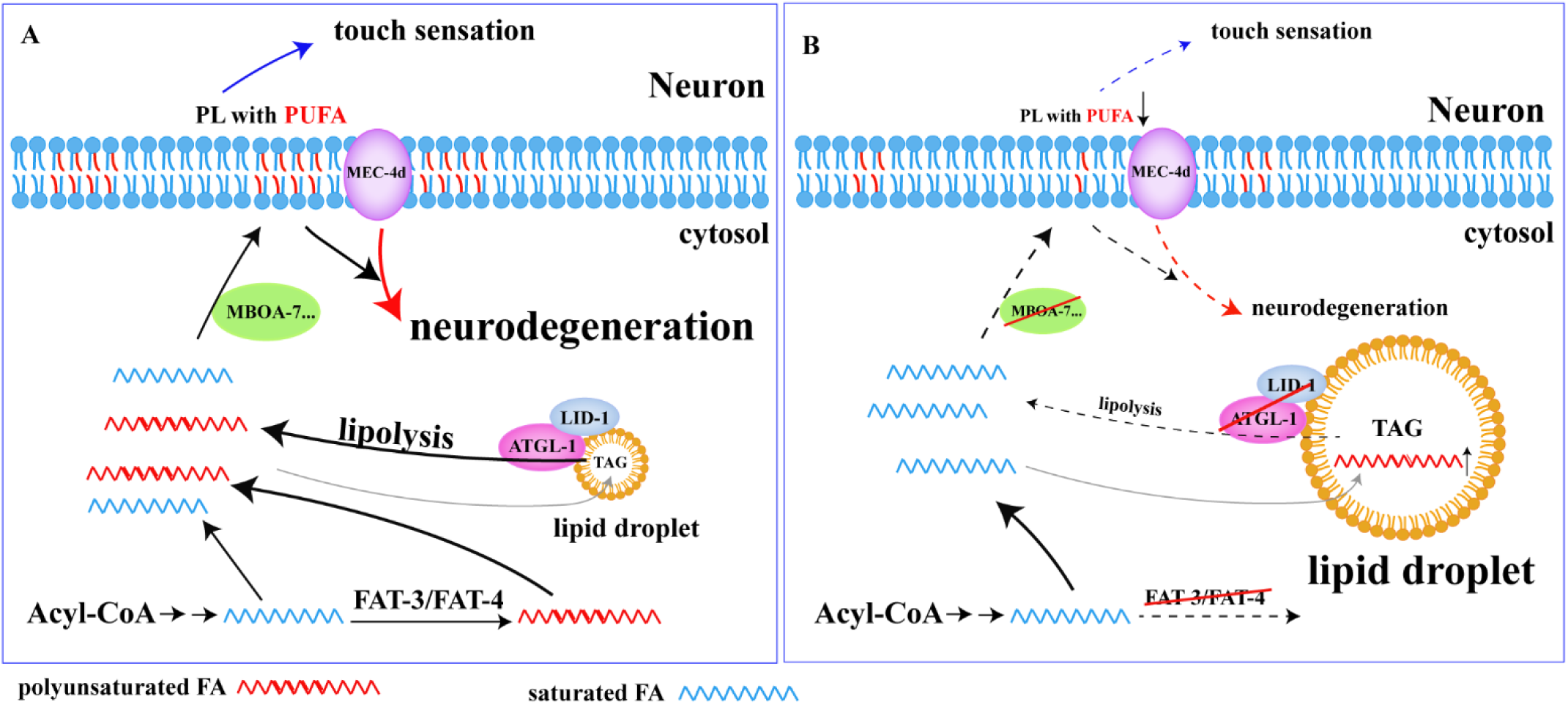
Neuronal lipolysis participates in the homeostasis of PUFAs, which regulates neuronal functions and neurodegeneration through incorporation into phospholipids. The homeostasis of PUFAs is determined by PUFA *de novo* synthesis with the balance of neuronal lipolysis and lipogenesis. The incorporation of PUFAs into phospholipids regulates neuronal normal functions (such as touch sensation) and affects neurodegeneration of *mec-4(d)*.

### LDs in neuron: a tug-of-war between lipogenesis and lipolysis

Neurons mainly use glucose to generate energy instead of lipids to avoid extra ROS production by lipid β-oxidation. Although normally there are no LDs in neurons, our results demonstrate that neurons have the ability to form and hydrolyze LDs. Both overexpression of lipogenesis genes and loss of the key lipolysis genes *atgl-1* and *lid-1* lead to LD accumulation in neurons. It is possible that the lipogenesis in neurons is limited and lipolysis is kept at a relatively high level, which promotes a fast turnover of LDs to meet the lipid requirements of neurons. Thus, there is a tug-of-war between lipogenesis and lipolysis in neurons.

Both *atgl-1* and *lid-1* are highly conserved from worm to mammals. Previous studies on *Drosophila* and cultured hippocampal neurons show that instead of forming LDs, neurons transfer the excess lipids into glial cells to form LDs under stress conditions (Bailey et al., 2015, Ioannou et al., 2019, Liu et al., 2017, Liu et al., 2015). If neurons have the ability to form and hydrolyze LDs, it is puzzling why they do not form LDs themselves. Could the formation of LDs in neurons be worm-specific because most neurons in *C. elegans* are not surrounded by glial cells? The appearance of LDs in *DDHD2*^*-/-*^ mice, which have a defect in a lipase, strongly argues against this possibility (Inloes, Hsu et al., 2014). Alternatively, it is possible that under conditions of intact neuronal lipolysis, excess lipids can still engage in generating toxic hyperoxidated PUFAs in neurons. Therefore, to avoid neuronal toxicity, exporting excess lipids to glia is a better choice. Examining the flux of lipid export and lipid incorporation into neutral lipids may provide a further explanation.

Interestingly, mice with deficiencies of *ATGL* or *CGI-58* accumulate massive amounts of neutral lipids in many tissues/organs including brain (Etschmaier, Becker et al., 2011, Radner, Streith et al., 2010). Similarly, lipid accumulation in brain was clearly demonstrated in *CGI-58* human patients (Huigen, van der Graaf et al., 2015). However, in both mouse and human studies, there is no direct evidence of LD accumulation in neurons. Compared to LDs in glia or blood vessels, neuronal LDs may be too small to observe and may therefore be neglected. Specific examination of neuronal LDs in these lipolysis-defective mutants will be required to characterize the neuronal defects.

### Neuronal lipolysis and neuronal normal function

Here, we show that intact neuronal lipolysis is important for maintaining normal neuron function in worm. Currently, there is no study on neurological function of ATGL-mediated lipolysis in mice. In patients with mutated *CGI-58*, neurological symptoms, including cognitive impairment and psychiatric disorders, are a frequent characteristic (Schweiger et al., 2009). A human *ATGL* mutation case with global cognitive impairment was also reported (Massa et al., 2016). Therefore, the critical role of neuronal lipolysis in neurons is likely conserved. The detail neuron-specific pathological phenotype and its underlying mechanism remain to be elucidated in *ATGL* or *CGI-58* mouse mutant models and more importantly in human patients.

Our results show that neuronal lipolysis participates in PUFA-mediated touch sensation. Through incorporation into membrane-forming phospholipids or as signals regulating neuron activity, PUFAs and their derivatives play important roles in neurons (Watts & Ristow, 2017). In *C. elegans* mechanical and touch sensation, PUFA-containing phospholipids modulate the activity of particular channels through membrane remodeling and changes of membrane fluidity (Matsuda, Inoue et al., 2008, Vasquez et al., 2014). In addition, PUFA depletion causes defects in neurotransmission (Lesa, Palfreyman et al., 2003, Marza & Lesa, 2006). It remains to be explored whether the neurological defects in *ATGL* or *CGI-58* human patients are linked to altered PUFA-mediated neuronal functions.

### Neuronal LDs and neurodegeneration

In this study, we found that LDs have a neuroprotective effect in the neuron hyperactivation-triggered neurodegeneration model. Interestingly, neuronal LDs also show a protective effect in another neurodegenerative disease, Parkinson’s disease. The expression of the Parkinson’s disease protein α-Synuclein alters the cellular lipid profile, notably the elevation of DAG and unsaturated fatty acids. By partitioning excess DAG and unsaturated fatty acids into TAG, the formation of LDs reduces neuron death triggered by expression of α-Synuclein (Fanning, Haque et al., 2019). Therefore, the presence of LDs may be beneficial for neurodegenerative diseases in general.

Besides Parkinson’s disease, many other neuronal diseases, in particular the HSP diseases, are reported to be accompanied by abnormal levels of neutral lipids, and the disease genes are related to LD dynamics. Knockdown of Spartin, also known as SPG20, increases the number and size of LDs in cells loaded with oleic acid (Papadopoulos et al., 2015). In another HSP model, *DDHD2*^*-/-*^ mice exhibit LD accumulation in neurons (Inloes et al., 2014). Despite the alteration of LD homeostasis, the role of LD accumulation in neurons under these disease conditions has not been investigated. Our findings also raise the possibility that the accumulation of LDs is a compensatory response to relieve neuronal stress in some HSP diseases. Further studies will be required to explore this possibility.

### PUFAs and neurodegeneration

We show that reducing PUFA incorporation into phospholipids has a beneficial effect in alleviating neuronal loss in a neurodegeneration model. The mechanism underlying this effect is not fully clear. Phospholipids are building blocks of all membranes in mammalian cells, providing the structural integrity that is necessary for protein function. They also serve as precursors for various second messengers such as AA, DHA, ceramide, 1,2-diacylglycerol, IP3, phosphatidic acid, and lyso-phospholipids (O’Donnell, Rossjohn et al., 2018). Generally, phospholipids with PUFA chains are more flexible. The cis-double bond lowers the packing density of the acyl chains, which increases membrane fluidity, bending stiffness and effective viscosity (Holthuis & Menon, 2014, Vasquez et al., 2014). One possibility is that a reduced level of PUFA-containing phospholipids impacts membrane fluidity and in turn reduces ion-channel activity. For example, AA and DHA can regulate channels such as K^+^ channels and TRPV4 (Caires, Sierra-Valdez et al., 2017, Horimoto, Nabekura et al., 1997, Villarroel & Schwarz, 1996). Moreover, phospholipids with PUFAs can be degraded and produce the second messengers 2-AG, 1,2-diacylglycerol and IP3. Therefore, another possibility is that reduced levels of phospholipids with PUFAs affect downstream signals that execute neurodegeneration triggered by ion-channel hyperactivation.

Our study indicates that reducing the influx of PUFAs into neuron phospholipids protects neurons from degeneration triggered by hyperactivation. This suggests a potential strategy for treating neurodegeneration caused by ion hyperactivation.

## Materials and Methods

### Strains

Worms were cultured on OP50-seeded nematode growth medium (NGM) plates at 22°C (Brenner, 1974). Wild-type (N2), CB4856, VC3025 *C37H5.2(ok3722)*, RB2386 *C37H5.3(ok3245)*, TU253 *mec-4(u253)*, CB1370 *daf-2(e1370)*, CB4037 *glp-1(e2141)*, CB1611 *mec-4(e1611)*, VC942 *mboa-7(gk399)* and GA184 *sod-2(gk257)* were obtained from the *Caenorhabditis* Genetics Center (CGC). *B0524.2(tm6739)* and *D1054.1(tm3111)* were obtained from the NBRP (Japan). Mutant strains BX24 *fat-1(wa9)*, BX30 *fat-3(wa22)* and BX17 *fat-4(wa14)* were kindly provided by Dr. Bin Liang. *lid-1(xd288), atgl-1(xd310)* and *atgl-1(xd314)* were generated by EMS. *xdIs182 (Prab-3::Cre/Podr-1::RFP)* was generated by integration of *Ex(Prab-3::Cre/Podr-1::RFP). atgl-1-loxP(xd426)* was generated by CRISPR-Cas9-mediated genome editing. All the transgenic worms were generated by microinjection of the respective plasmids or fosmids with co-injection markers. All the fosmids were kindly provided by Dr. Xiaochen Wang.

### Molecular biology

The promoters of P*unc-119* (1235bp), P*vha-6* (1808bp), P*hlh-17* (2049bp), P*ajm-1* (3500bp), P*rab-3* (1207bp), P*atgl-1* (3004bp) and P*mec-4* (206bp) were amplified from N2 genomic DNA. The coding regions of *lid-1* (1083bp), *atgl-1* (2256bp), *C37H5.2* (1068bp), *C37H5.3* (1335bp), *B0524.2* (1047bp), *D1054.1* (801bp) and *mboa-2* (1497bp) were amplified from cDNA. The promotor and target cDNA were inserted into plasmid *Δpsm*. The split GFP1-10 and GFP11 were inserted into *pPD95.75*. P*atgl-1* and P*rab-3* were inserted upstream of GFP1-10 and GFP11 respectively. *Prab-3::Cre* was generated by replacing the *eft-3* promotor in *pDD104*, which was kindly provided by Dr. Shiqing Cai. *atgl-1*-sgRNAs for loxP insertion were designed using the Zhang lab’s CRISPR design tool at *http://crispr.mit.edu* to select the target sites. The sgRNAs were inserted into *pDD162*, which was kindly provided by Dr. Guangshuo Ou. All the constructed plasmids were verified by sequencing.

### EMS screen

We treated L4 and young adult *xdIs109 (Punc-119::PLIN1::GFP)* worms with ethyl methanesulfonate (EMS). About 2,500 F1 were screened. We observed LDs using a Zeiss compound microscope. Mutants were selected that showed LD accumulation in neurons.

### SNP mapping and fosmid rescue

We used rapid Single Nucleotide Polymorphism (SNP) mapping (Davis, Hammarlund et al., 2005). Briefly, we crossed the mutant with CB4856 and picked F2 animals with LD accumulation. We amplified some SNPs using F2 or F3 lysates as templates and we digested the PCR products with enzymes. After chromosome mapping and interval mapping, we identified a narrow region. Then we did the fosmid rescue assay. Fosmids were injected individually into mutant worms at 5∼10 ng/μl together with 50 ng/μl *Podr-1::RFP*.

### Oil red O staining

Oil red O was acquired from Sigma-Aldrich. Oil red O staining was conducted as previously reported (O’Rourke et al., 2009).

### Behavioral assays

Gentle touch sensitivity was tested and scored as described (Hart, 2006, Vasquez et al., 2014). Briefly, we performed ten-trial touch assays. We scored the touch response percentage by stroking an eyebrow across the anterior and posterior body. Twenty-five animals were tested in each trial, and results were compared across three trials. All assays were performed by investigators blinded to genotype and/or treatment.

### Quantification of neuron loss

Animals were mounted on thin agarose pads and immobilized by 1-phenoxy-2-propanol (10 μL/mL). Animals were visualized under a 20X objective. GFP-expressing touch neurons were counted in synchronized L4 animals.

### Fatty acid and NAC supplementation

AA (arachidonic acid), and NAC (N-Acetyl-L-cysteine) were acquired from Aladdin. SA (stearic acid), OA (oleic acid), PA (palmitate acid), EPA (eicosapentaenoic acid), LA (linoleic acid) and Tergitol (70%) were acquired from Sigma-Aldrich. FAs, dissolved in ethanol with 0.1% Tergitol, were added to NGM agar to reach a final concentration of approximately 200 mM. NAC was dissolved in ddH_2_O and added into NGM agar to reach a final concentration of approximately 10 mM. Plates were seeded with *E. coli* OP50 and kept at room temperature (Vasquez et al., 2014). For each strain, about 10 L4 worms were placed on the plate. We quantified the phenotype in the next generation.

### Stimulated Raman Scattering (SRS) microscopy

To quantify lipid droplet metrics in the touch neurons, 1-day adult worms were imaged using Stimulated Raman Scattering (SRS) microscopy. Worms were mounted onto 2% agarose pads with 0.5% NaN3 as anesthetic on glass microscope slides and imaged using a 60× water objective (UPlanAPO/IR; 1.2 N.A.; Olympus). A femtosecond-pulsed laser and picosecond-pulsed laser were used for simultaneously imaging label-free lipids (SRS channel) and GFP-labeled neurons (fluorescence channel). Images were analyzed using ImageJ software (NIH). Neuronal area was selected using fluorescent signal based on several Z-projected stacks to confirm the presence of lipid droplets within the cell and to exclude extracellular signal.

### EM analysis

Young adult worms were collected for high-pressure freezing and freeze-substitution, embedding and sectioning following a procedure essentially as described by (Weimer, 2006). Then the sections were visualized with a Hitachi HT7700 and pictures were recorded on a 4008 × 2672 CCD camera.

### Lipidomic analysis

Worms were washed 9∼10 times in M9. More than 10,000 worms per genotype were collected per sample and 5 samples were analyzed per genotype. Lipid extraction and analysis were conducted as previously reported (Lam, Wang et al., 2014). The lipid content was normalized by the mole fraction of each lipid to total polar lipids.

### Statistical analyses

All data are presented as mean ± SEM. The data were first analyzed using one-way ANOVA or two-way ANOVA and for statistically significant overall results. Post hoc multiple comparison tests were carried out to determine significant differences. Significant difference is noted with a number sign (#) or an asterisk (*). ns represents not statistically significant. *\*P* < 0.05, *\*\*P* < 0.01, *\*\*\*P* < 0.001, \**\*\*\*P* < 0.0001. #*P* < 0.05, ##*P* < 0.01, ###*P* < 0.001, ####*P* < 0.0001.

## Acknowledgements

We thank Drs. Shiqing Cai, Bin Liang, Guangshuo Ou and Xiaochen Wang for providing reagents and helpful discussions. We thank CGC and NBRP for providing strains. This research was supported by grants 31630019/9195420001 and 2016YFA0500100/2018YFA0506902 from the National Natural Science Foundation of China and the Ministry of Science and Technology of China, respectively.

## Author contributions

Y. L. directed the project and her work includes EMS screen, molecular biology, SNP mapping and fosmid rescue, Oil red O staining, behavioral assays, neuronal loss analysis, fatty acid and NAC supplementation and analyzing the results. L. J. and D. M. did the EM imaging and analysis. L. S. M. and S. G. did the lipidomic analysis. Y. A. and W. C. M did the SRS imaging. Y. L. and H. X. wrote the manuscript. H. X. guided the project and edited the manuscript.

---

**Fig. S1.**
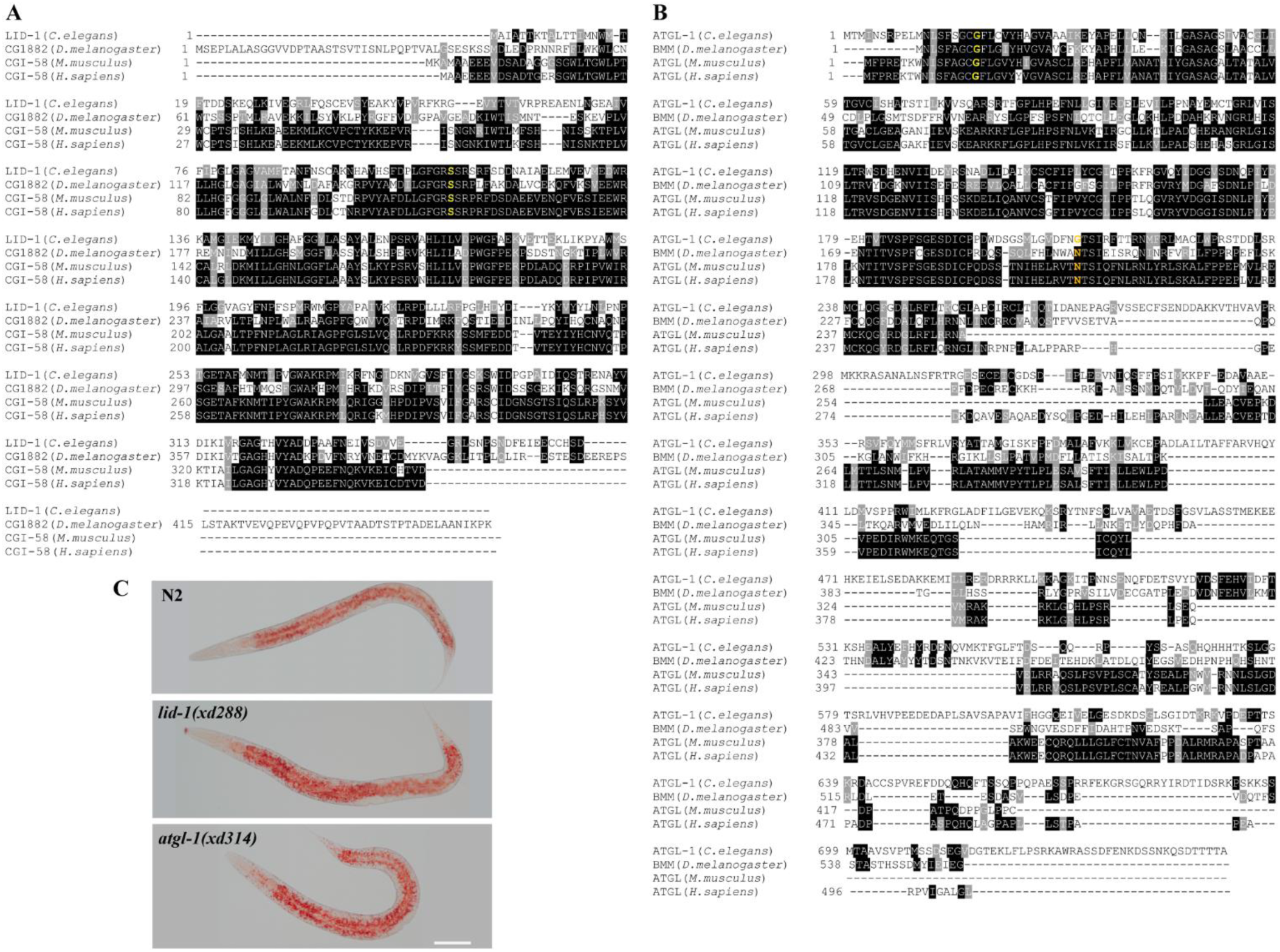
LID-1 and ATGL-1 are conserved. (A) and (B) Alignment of CGI-58 and ATGL from *C. elegans, D. melanogaster, M. musculus* and *H. sapiens*. Single-letter abbreviations for amino acid residues are used. Identical and similar amino acids are identified by gray and light gray shading, respectively. The yellow and orange letters indicate the point mutation sites. (C) Oil red O staining (red) shows that the overall neutral lipids are increased in *lid-1(xd288)* and *atgl-1(xd314)*. The worms are at L4 stage. Scale bar: 50 μm.

**Fig. S2.**
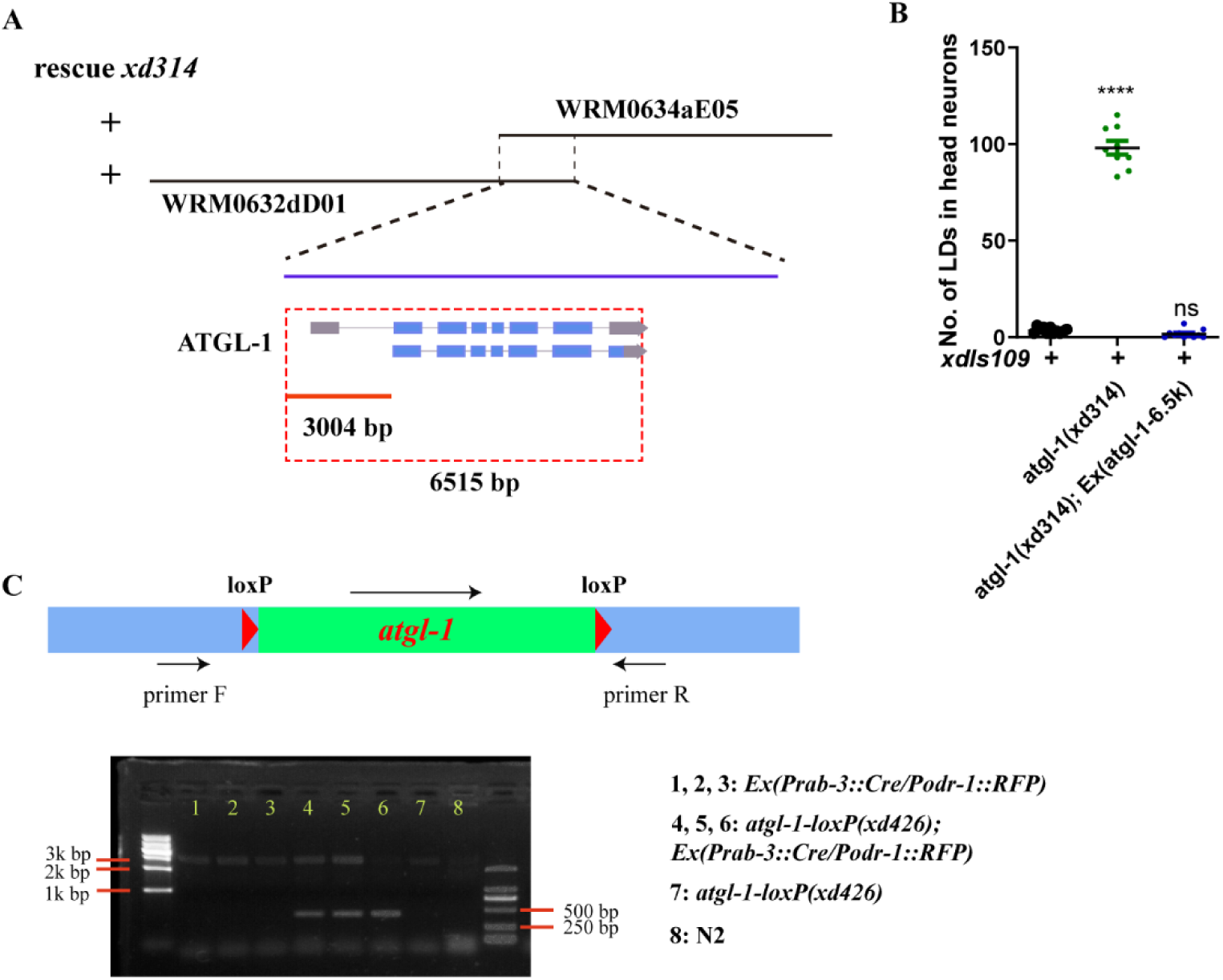
*Patgl-1::atgl-1* fully rescues the phenotype of neuronal LD accumulation in *atgl-1(xd314)* mutants. (A) The rescue of *atgl-1(xd314)* by both WRM0632dD01 and WRM0634aE05 fosmids narrows down the functional promoter of *atgl-1* to 3,400 bp upstream of the start codon ATG. (B) Quantification of the number of LDs in head neurons in different genetic backgrounds. The 6,515 bp *atgl-1(+)* rescues the phenotype of *atgl-1(xd314)*. Each dot represents one worm. The data was analyzed using one-way ANOVA with Dunnett’s multiple comparison test. * signifies the significant difference as compared to control *xdIs109*. Error bars, SEM. n ≥ 9. ns not statistically significant. (C) LoxP were inserted into the two sides of *atgl-1* locus. To identify *atgl-1* deletions, primer F and primer R were used to amplify *atgl-1* locus. There is only one band in negative control *Ex(Prab-3::Cre/Podr-1::RFP)* and *atgl-1-loxP(xd426)*. There are two bands (one large and one small) in *atgl-1-loxP(xd426); Ex(Prab-3::Cre/Podr-1::RFP)*, indicating that *atgl-1* is partially deleted.

**Fig. S3.**
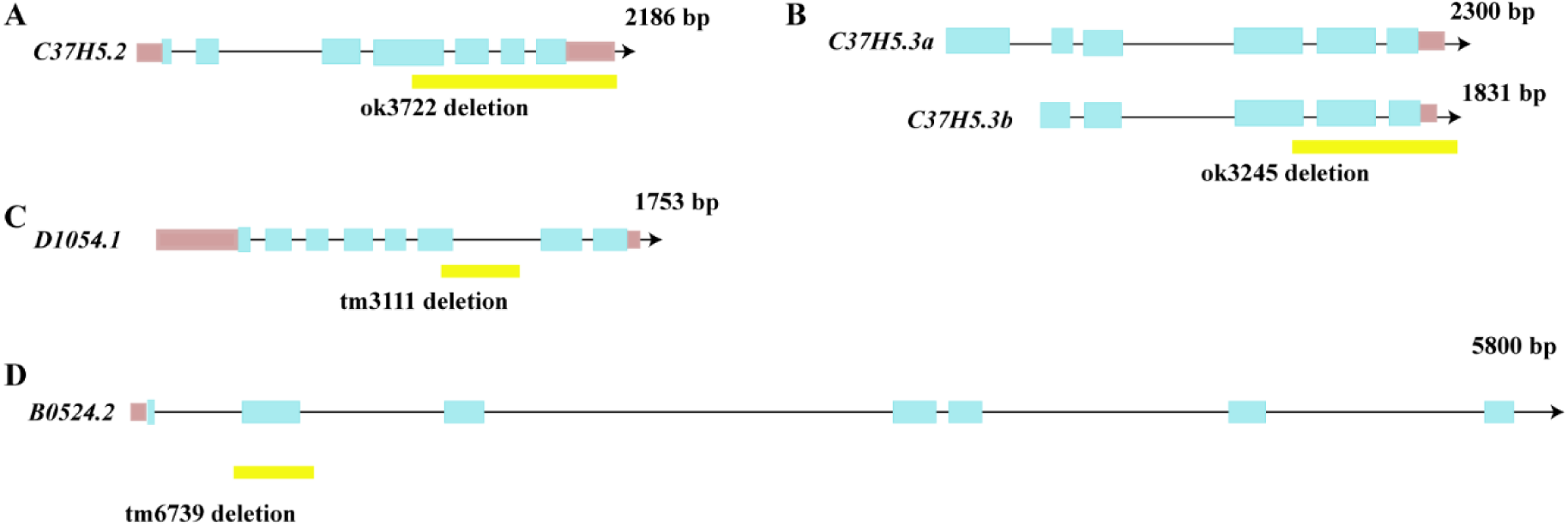
The schematic diagram of deletion region in mutants of LID-1/ATGL-1 paralogs. (A) to (D), the schematic diagram of alleles of LID-1/ATGL-1 paralogs. The coding regions are in blue boxes and the noncoding regions are shown as lines. The UTRs are in pink boxes. The yellow boxes show the regions deleted in the corresponding alleles.

**Fig. S4.**
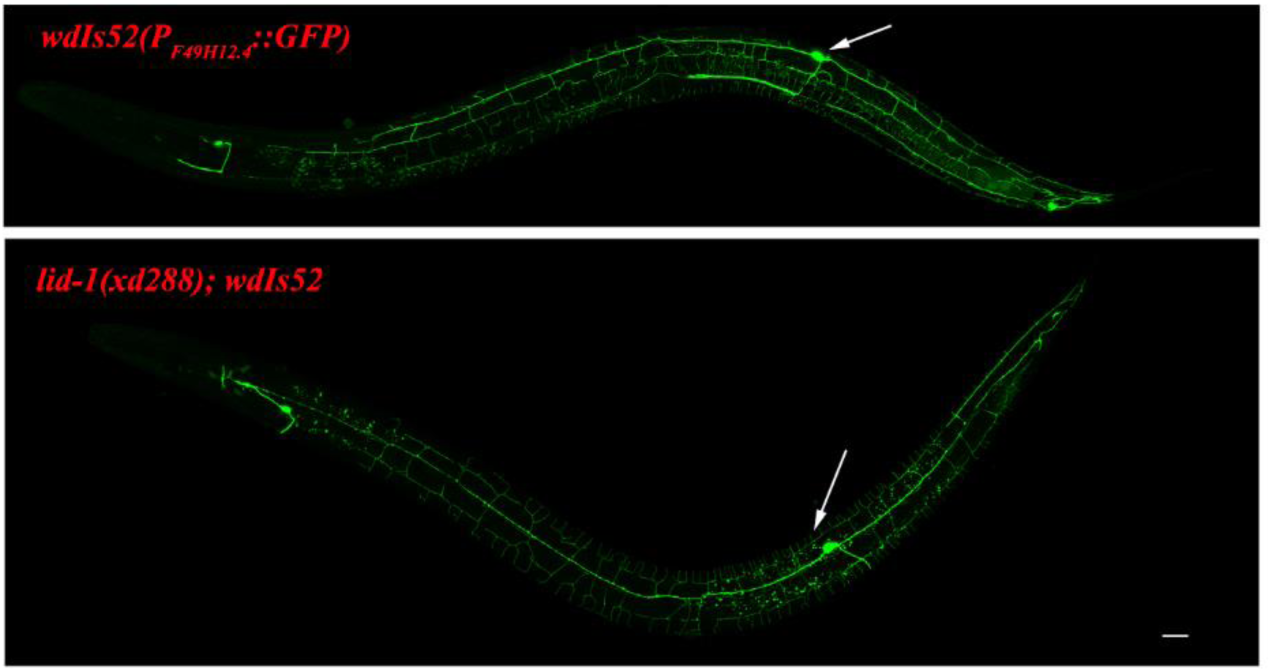
The mutation of *lid-1* doesn’t affect the morphology of PVD. *P*_*F49H12.4*_::*GFP* reporter marks the morphology of neurons PVDL/R. The white arrows indicate the soma of PVD. The neurites of PVD almost cover the whole worm body. There is no obvious defect of PVD morphology in *lid-1(xd288)* viewed by *wdIs52(P*_*F49H12.4*_::*GFP)*. Scale bar: 20 μm.

**Fig. S5.**
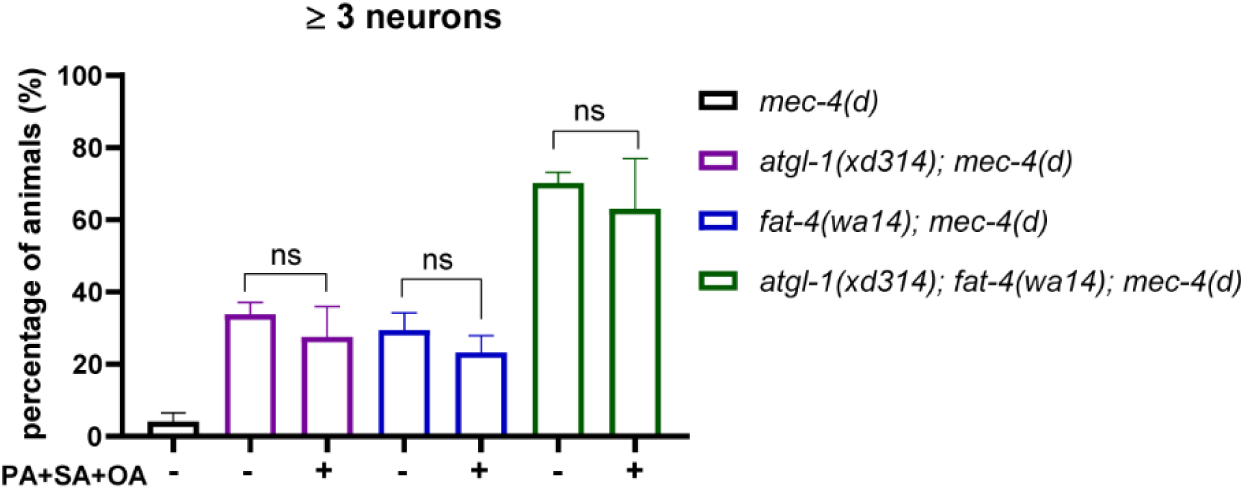
Saturated fatty acids and mono-unsaturated fatty acids don’t significantly affect neurodegeneration triggered by *mec-4(d)*. The percentage of worms that have three or more surviving touch neurons in different genetic backgrounds supplemented with fatty acid mix. The supplement of PA, SA and OA don’t affect the neurodegeneration triggered by *mec-4(d)*. The data was analyzed using one-way ANOVA with Bonferroni’s multiple comparison test. ns: not statistically significant. Error bars, SEM. Number of experiments n ≥ 3, with at least 50 animals per strain analyzed in each experiment.

**Fig. S6.**
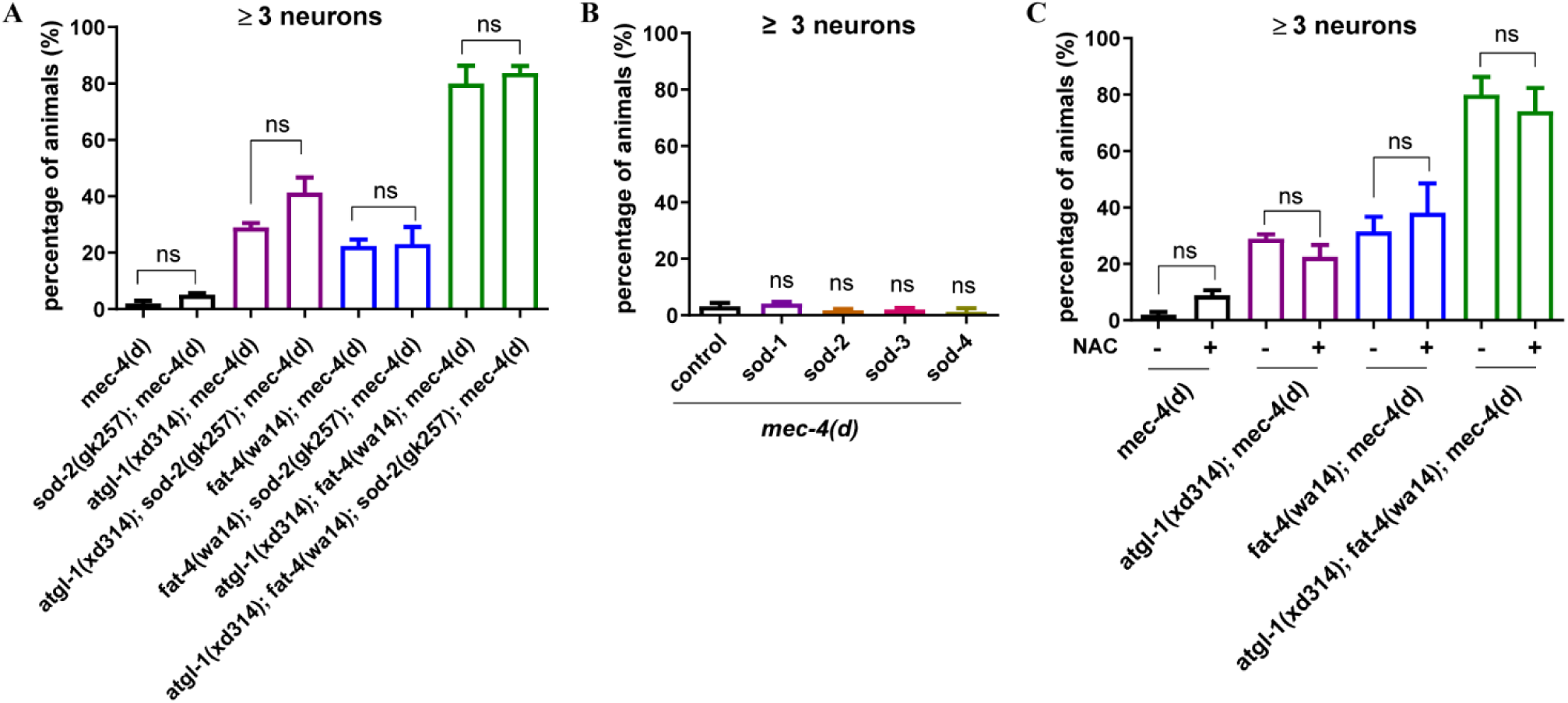
Manipulating ROS scavengers doesn’t influence protective effect of lipolysis mutants or PUFA mutants in neurodegeneration caused by *mec-4(d)* (A) The percentage of worms that have three or more surviving touch neurons in different genetic backgrounds. *sod-2* mutation doesn’t enhance neurodegeneration in *mec-4(d), atgl-1(xd314); mec-4(d), fat-4(wa14); mec-4(d)* and *atgl-1(xd314); fat-4(wa14); mec-4(d)*. The data was analyzed using one-way ANOVA with Bonferroni’s multiple comparison test. Error bars, SEM. ns: not statistically significant. Number of experiments n ≥ 5, with at least 50 animals per strain analyzed in each experiment. (B) The percentage of worms that have three or more surviving touch neurons in different genetic backgrounds. Using touch neuron specific promoter *Pmec-4* driving *sod-1, sod-2, sod-3* or *sod-4* overexpression in touch neurons doesn’t significantly affect neurodegeneration in *mec-4(d)*. The data was analyzed using one-way ANOVA with Bonferroni’s multiple comparison test. Error bars, SEM. ns: not statistically significant. Number of experiments n ≥ 2, with at least 50 animals per strain analyzed in each experiment. (C) The percentage of worms that have three or more surviving touch neurons in different genetic backgrounds. Feeding worms with antioxidant NAC doesn’t significantly affect neurodegeneration in *mec-4(d), atgl-1(xd314); mec-4(d), fat-4(wa14); mec-4(d)* and *atgl-1(xd314); fat-4(wa14); mec-4(d)*. The data was analyzed using one-way ANOVA with Bonferroni’s multiple comparison test. Error bars, SEM. ns: not statistically significant. Number of experiments n ≥ 5, with at least 50 animals per strain analyzed in each experiment.

**Fig. S7.**
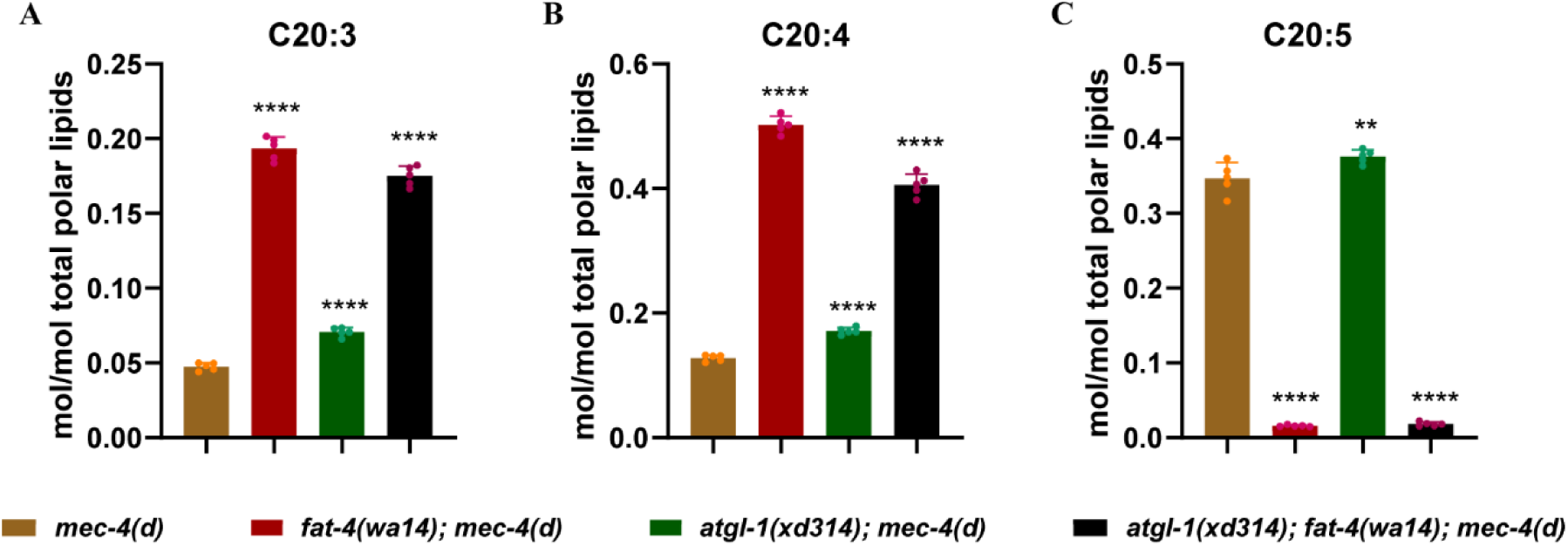
The relative content of lipids that have C20:3, C20:4 or C20:5 PUFAs. (A) Lipidomic data show that the relative content of total C20:3 is increased in *atgl-1(xd314); mec-4(d), fat-4(wa14); mec-4(d)* and *atgl-1(xd314); fat-4(wa14); mec-4(d)*, especially in mutants that have *fat-4* mutation. (B) Lipidomic data show that the relative content of total C20:4 is increased in *atgl-1(xd314); mec-4(d), fat-4(wa14); mec-4(d)* and *atgl-1(xd314); fat-4(wa14); mec-4(d)*, especially in mutants that have *fat-4* mutation. (C) Lipidomic data show that the relative content of total C20:5 is slightly increased in *atgl-1(xd314); mec-4(d)* but a dramatically decreased in *fat-4(wa14); mec-4(d)* and *atgl-1(xd314); fat-4(wa14); mec-4(d)*, which matches the role of FAT-4 in synthesizing C20:5 (EPA). (A-C) The data was analyzed using one-way ANOVA with Dunnett’s multiple comparison test. * signifies the significant difference as compared to control *mec-4(d)*. Error bars, SEM. n=5.

**Fig. S8.**
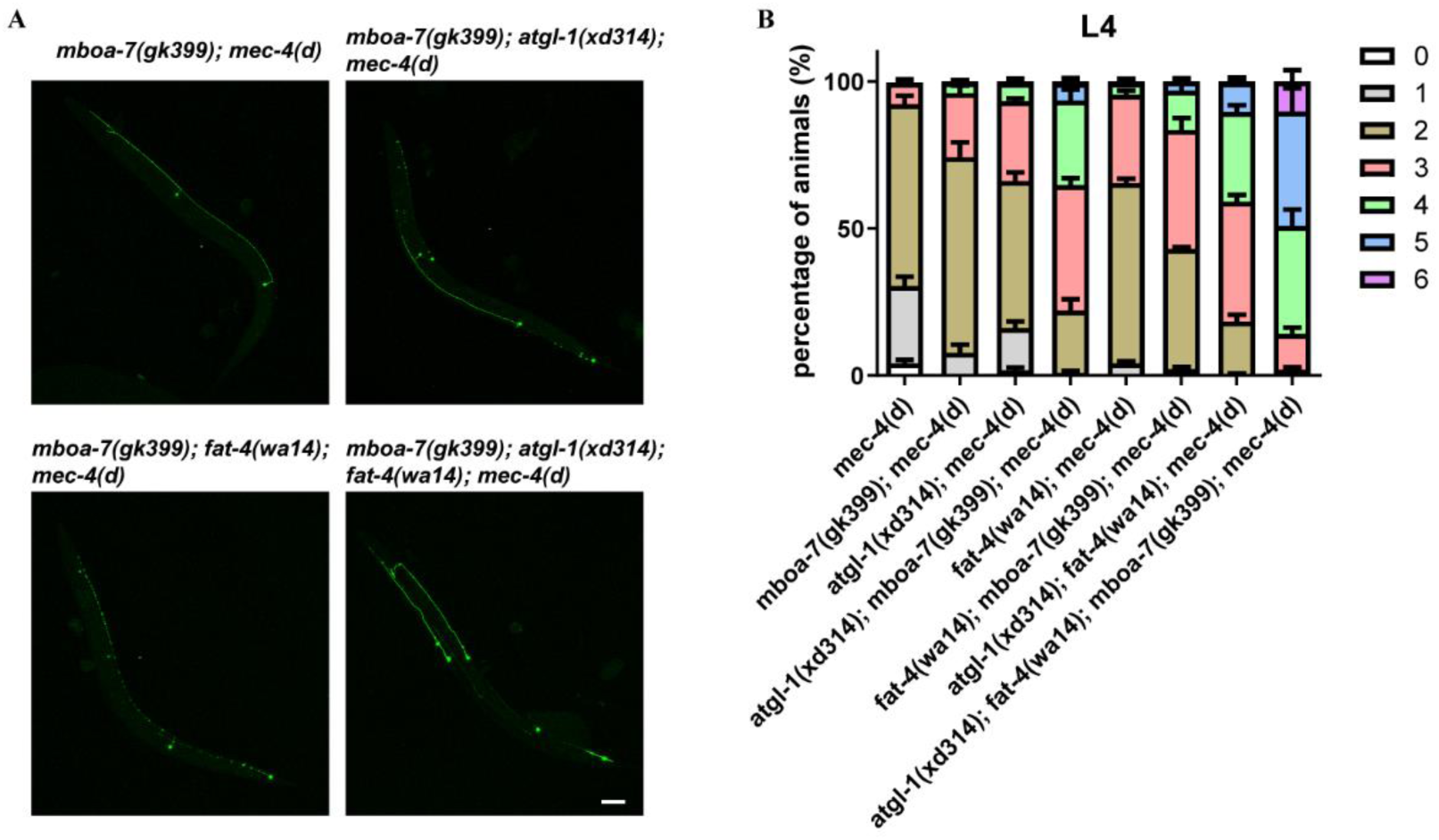
*mboa-7* mutation significantly affects neurodegeneration caused by *mec-4(d)*. (A) Image of touch neurons in worms viewed by *zdIs5(Pmec-4::GFP)* in different genetic backgrounds. Scale bar: 20 μm. (B) Quantification of the percentage of animals with 0, 1, 2, 3, 4, 5 and 6 surviving touch neurons in different genetic backgrounds. The distribution of surviving neuron number is further improved when *mboa-7* is mutated in *atgl-1, fat-4* single mutants or double mutant. At least 50 animals per strain were analyzed in each experiment. Number of experiments was 5.

## References

Bailey AP, Koster G, Guillermier C, Hirst EM, MacRae JI, Lechene CP, Postle AD, Gould AP (2015) Antioxidant role for lipid droplets in a stem cell niche of *Drosophila*. Cell 163: 340–53

Ben Selma Z, Yilmaz S, Schischmanoff PO, Blom A, Ozogul C, Laroche L, Caux F (2007) A novel S115G mutation of CGI-58 in a Turkish patient with Dorfman-Chanarin syndrome. The Journal of investigative dermatology 127: 2273–6

Bi J, Xiang Y, Chen H, Liu Z, Gronke S, Kuhnlein RP, Huang X (2012) Opposite and redundant roles of the two *Drosophila* perilipins in lipid mobilization. J Cell Sci 125: 3568–77

Brenner S (1974) The genetics of *Caenorhabditis elegans*. Genetics 77: 71–94

Cabantous S, Terwilliger TC, Waldo GS (2005) Protein tagging and detection with engineered self-assembling fragments of green fluorescent protein. Nat Biotechnol 23: 102–7

Caires R, Sierra-Valdez FJ, Millet JRM, Herwig JD, Roan E, Vasquez V, Cordero-Morales JF (2017) Omega-3 fatty acids modulate TRPV4 function through plasma membrane remodeling. Cell Rep 21: 246–258

Calixto A, Jara JS, Court FA (2012) Diapause formation and downregulation of insulin-like signaling via DAF-16/FOXO delays axonal degeneration and neuronal loss. PLoS Genet 8: e1003141

Chen L, Chen X-W, Huang X, Song B-L, Wang Y, Wang Y (2019) Regulation of glucose and lipid metabolism in health and disease. Sci China Life Sci: doi: 10.1007/s11427-019-1563-3

Cole NB, Murphy DD, Grider T, Rueter S, Brasaemle D, Nussbaum RL (2002) Lipid droplet binding and oligomerization properties of the Parkinson’s disease protein alpha-synuclein. J Biol Chem 277: 6344–52

Davis MW, Hammarlund M, Harrach T, Hullett P, Olsen S, Jorgensen EM (2005) Rapid single nucleotide polymorphism mapping in *C. elegans*. BMC Genomics 6: 118

Ding L, Yang X, Tian H, Liang J, Zhang F, Wang G, Wang Y, Ding M, Shui G, Huang X (2018) Seipin regulates lipid homeostasis by ensuring calcium-dependent mitochondrial metabolism. EMBO J 37: e97572

Driscoll M, Chalfie M (1991) The *mec-4* gene is a member of a family of *Caenorhabditis elegans* genes that can mutate to induce neuronal degeneration. Nature 349: 588–93

Eastman SW, Yassaee M, Bieniasz PD (2009) A role for ubiquitin ligases and Spartin/SPG20 in lipid droplet turnover. J Cell Biol 184: 881–94

Ebihara C, Ebihara K, Aizawa-Abe M, Mashimo T, Tomita T, Zhao M, Gumbilai V, Kusakabe T, Yamamoto Y, Aotani D, Yamamoto-Kataoka S, Sakai T, Hosoda K, Serikawa T, Nakao K (2015) Seipin is necessary for normal brain development and spermatogenesis in addition to adipogenesis. Hum Mol Genet 24: 4238–49

Etschmaier K, Becker T, Eichmann TO, Schweinzer C, Scholler M, Tam-Amersdorfer C, Poeckl M, Schuligoi R, Kober A, Chirackal Manavalan AP, Rechberger GN, Streith IE, Zechner R, Zimmermann R, Panzenboeck U (2011) Adipose triglyceride lipase affects triacylglycerol metabolism at brain barriers. J Neurochem 119: 1016–28

Fanning S, Haque A, Imberdis T, Baru V, Barrasa MI, Nuber S, Termine D, Ramalingam N, Ho GPH, Noble T, Sandoe J, Lou Y, Landgraf D, Freyzon Y, Newby G, Soldner F, Terry-Kantor E, Kim TE, Hofbauer HF, Becuwe M et al. (2019) Lipidomic analysis of alpha-synuclein neurotoxicity identifies stearoyl CoA desaturase as a target for parkinson treatment. Mol Cell 73: 1001–1014 e8

Hart AC (2006) Behavior. WormBook, doi/10.1895/wormbook.1.87.1

Holthuis JC, Menon AK (2014) Lipid landscapes and pipelines in membrane homeostasis. Nature 510: 48–57

Horimoto N, Nabekura J, Ogawa T (1997) Arachidonic acid activation of potassium channels in rat visual cortex neurons. Neuroscience 77: 661–71

Huigen MC, van der Graaf M, Morava E, Dassel AC, van Steensel MA, Seyger MM, Wevers RA, Willemsen MA (2015) Cerebral lipid accumulation in Chanarin-Dorfman Syndrome. Mol Genet Metab 114: 51–4

Inloes JM, Hsu KL, Dix MM, Viader A, Masuda K, Takei T, Wood MR, Cravatt BF (2014) The hereditary spastic paraplegia-related enzyme DDHD2 is a principal brain triglyceride lipase. Proc Natl Acad Sci USA 111: 14924–9

Ioannou MS, Jackson J, Sheu SH, Chang CL, Weigel AV, Liu H, Pasolli HA, Xu CS, Pang S, Matthies D, Hess HF, Lippincott-Schwartz J, Liu Z (2019) Neuron-astrocyte metabolic coupling protects against activity-induced fatty acid toxicity. Cell 177: 1522–1535 e14

Kahn-Kirby AH, Dantzker JL, Apicella AJ, Schafer WR, Browse J, Bargmann CI, Watts JL (2004) Specific polyunsaturated fatty acids drive TRPV-dependent sensory signaling *in vivo*. Cell 119: 889–900

Kis V, Barti B, Lippai M, Sass M (2015) Specialized cortex glial cells accumulate lipid droplets in *Drosophila melanogaster*. PLoS One 10: e0131250

Klemm RW, Norton JP, Cole RA, Li CS, Park SH, Crane MM, Li L, Jin D, Boye-Doe A, Liu TY, Shibata Y, Lu H, Rapoport TA, Farese RV, Jr., Blackstone C, Guo Y, Mak HY (2013) A conserved role for atlastin GTPases in regulating lipid droplet size. Cell Rep 3: 1465–75

Lam SM, Wang Y, Duan X, Wenk MR, Kalaria RN, Chen CP, Lai MK, Shui G (2014) Brain lipidomes of subcortical ischemic vascular dementia and mixed dementia. Neurobiol Aging 35: 2369–81

Lass A, Zimmermann R, Haemmerle G, Riederer M, Schoiswohl G, Schweiger M, Kienesberger P, Strauss JG, Gorkiewicz G, Zechner R (2006) Adipose triglyceride lipase-mediated lipolysis of cellular fat stores is activated by CGI-58 and defective in Chanarin-Dorfman Syndrome. Cell metabolism 3: 309–19

Lee HC, Inoue T, Imae R, Kono N, Shirae S, Matsuda S, Gengyo-Ando K, Mitani S, Arai H (2008) *Caenorhabditis elegans mboa-7*, a member of the MBOAT family, is required for selective incorporation of polyunsaturated fatty acids into phosphatidylinositol. Mol Biol Cell 19: 1174–84

Lee JH, Kong J, Jang JY, Han JS, Ji Y, Lee J, Kim JB (2014) Lipid droplet protein LID-1 mediates ATGL-1-dependent lipolysis during fasting in *Caenorhabditis elegans*. Mol Cell Biol 34: 4165–76

Lesa GM, Palfreyman M, Hall DH, Clandinin MT, Rudolph C, Jorgensen EM, Schiavo G (2003) Long chain polyunsaturated fatty acids are required for efficient neurotransmission in *C. elegans*. J Cell Sci 116: 4965–75

Liu L, MacKenzie KR, Putluri N, Maletic-Savatic M, Bellen HJ (2017) The glia-neuron lactate shuttle and elevated ROS promote lipid synthesis in neurons and lipid droplet accumulation in glia via APOE/D. Cell Metab 26: 719–737 e6

Liu L, Zhang K, Sandoval H, Yamamoto S, Jaiswal M, Sanz E, Li Z, Hui J, Graham BH, Quintana A, Bellen HJ (2015) Glial lipid droplets and ROS induced by mitochondrial defects promote neurodegeneration. Cell 160: 177–90

Liu Z, Li X, Ge Q, Ding M, Huang X (2014) A lipid droplet-associated GFP reporter-based screen identifies new fat storage regulators in C. elegans. J Genet Genomics 41: 305–13

Martinez-Vicente M, Talloczy Z, Wong E, Tang G, Koga H, Kaushik S, de Vries R, Arias E, Harris S, Sulzer D, Cuervo AM (2010) Cargo recognition failure is responsible for inefficient autophagy in Huntington’s disease. Nat Neurosci 13: 567–76

Marza E, Lesa GM (2006) Polyunsaturated fatty acids and neurotransmission in Caenorhabditis elegans. Biochem Soc Trans 34: 77–80

Massa R, Pozzessere S, Rastelli E, Serra L, Terracciano C, Gibellini M, Bozzali M, Arca M (2016) Neutral lipid-storage disease with myopathy and extended phenotype with novel PNPLA2 mutation. Muscle Nerve 53: 644–8

Matsuda S, Inoue T, Lee HC, Kono N, Tanaka F, Gengyo-Ando K, Mitani S, Arai H (2008) Member of the membrane-bound O-acyltransferase (MBOAT) family encodes a lysophospholipid acyltransferase with broad substrate specificity. Genes Cells 13: 879–88

Narbonne P, Roy R (2009) *Caenorhabditis elegans* dauers need LKB1/AMPK to ration lipid reserves and ensure long-term survival. Nature 457: 210–4

O’Donnell VB, Rossjohn J, Wakelam MJ (2018) Phospholipid signaling in innate immune cells. J Clin Invest 128: 2670–2679

O’Rourke EJ, Soukas AA, Carr CE, Ruvkun G (2009) *C. elegans* major fats are stored in vesicles distinct from lysosome-related organelles. Cell Metab 10: 430–5

Olzmann JA, Carvalho P (2019) Dynamics and functions of lipid droplets. Nat Rev Mol Cell Biol 20: 137–155

Outeiro TF, Lindquist S (2003) Yeast cells provide insight into alpha-synuclein biology and pathobiology. Science 302: 1772–5

Palikaras K, Mari M, Petanidou B, Pasparaki A, Filippidis G, Tavernarakis N (2017) Ectopic fat deposition contributes to age-associated pathology in *Caenorhabditis elegans*. J Lipid Res 58: 72–80

Papadopoulos C, Orso G, Mancuso G, Herholz M, Gumeni S, Tadepalle N, Jungst C, Tzschichholz A, Schauss A, Honing S, Trifunovic A, Daga A, Rugarli EI (2015) Spastin binds to lipid droplets and affects lipid metabolism. PLoS Genet 11: e1005149

Radner FP, Streith IE, Schoiswohl G, Schweiger M, Kumari M, Eichmann TO, Rechberger G, Koefeler HC, Eder S, Schauer S, Theussl HC, Preiss-Landl K, Lass A, Zimmermann R, Hoefler G, Zechner R, Haemmerle G (2010) Growth retardation, impaired triacylglycerol catabolism, hepatic steatosis, and lethal skin barrier defect in mice lacking comparative gene identification-58 (CGI-58). J Biol Chem 285: 7300–11

Renvoise B, Malone B, Falgairolle M, Munasinghe J, Stadler J, Sibilla C, Park SH, Blackstone C (2016) Reep1 null mice reveal a converging role for hereditary spastic paraplegia proteins in lipid droplet regulation. Hum Mol Genet 25: 5111–5125

Sangaletti R, D’Amico M, Grant J, Della-Morte D, Bianchi L (2017) Knock-out of a mitochondrial sirtuin protects neurons from degeneration in *Caenorhabditis elegans*. PLoS Genet 13: e1006965

Schweiger M, Lass A, Zimmermann R, Eichmann TO, Zechner R (2009) Neutral lipid storage disease: genetic disorders caused by mutations in adipose triglyceride lipase/PNPLA2 or CGI-58/ABHD5. Am J Physiol Endocrinol Metab 297: E289–96

Shimabukuro MK, Langhi LG, Cordeiro I, Brito JM, Batista CM, Mattson MP, Mello Coelho V (2016) Lipid-laden cells differentially distributed in the aging brain are functionally active and correspond to distinct phenotypes. Sci Rep 6: 23795

Vasquez V, Krieg M, Lockhead D, Goodman MB (2014) Phospholipids that contain polyunsaturated fatty acids enhance neuronal cell mechanics and touch sensation. Cell Rep 6: 70–80

Villarroel A, Schwarz TL (1996) Inhibition of the Kv4 (Shal) family of transient K+ currents by arachidonic acid. J Neurosci 16: 2522–32

Watts JL, Browse J (2002) Genetic dissection of polyunsaturated fatty acid synthesis in *Caenorhabditis elegans*. Proc Natl Acad Sci USA 99: 5854–9

Watts JL, Ristow M (2017) Lipid and carbohydrate metabolism in *Caenorhabditis elegans*. Genetics 207: 413–446

Weimer RM (2006) Preservation of *C. elegans* tissue via high-pressure freezing and freeze-substitution for ultrastructural analysis and immunocytochemistry. Methods Mol Biol 351: 203–21

Xie M, Roy R (2015a) AMP-Activated Kinase Regulates Lipid Droplet Localization and Stability of Adipose Triglyceride Lipase in *C. elegans* Dauer Larvae. PLoS One 10: e0130480

Xie M, Roy R (2015b) The causative gene in Chanarian Dorfman Syndrome regulates lipid droplet homeostasis in *C. elegans*. PLoS Genet 11: e1005284

Zhou YT, Grayburn P, Karim A, Shimabukuro M, Higa M, Baetens D, Orci L, Unger RH (2000) Lipotoxic heart disease in obese rats: implications for human obesity. Proc Natl Acad Sci USA 97: 1784–9

